# Single molecule dynamics of Dishevelled at the plasma membrane and Wnt pathway activation

**DOI:** 10.1101/624882

**Authors:** Wenzhe Ma, Maorong Chen, Hong Kang, Zachary Steinhart, Stephane Angers, Xi He, Marc W. Kirschner

**Affiliations:** Department of Systems Biology, Harvard Medical School, Boston, MA 02115, USA; F. M. Kirby Neurobiology Center, Children’s Hospital Boston, Harvard Medical School, Boston, MA 02115, USA; University of Toronto, 144 College Street, Toronto, ON M5S 3M2, Canada

**Keywords:** Wnt Signaling Pathway, Disheveled, Single Molecule, Protein Complex Size, Fluorescence, TIRF (Total Intensity Reflection Fluorescence), Kinetic Models of complex assembly

## Abstract

Dvl (Dishevelled) is one of several essential non-enzymatic components of the Wnt signaling pathway. In most current models, Dvl forms complexes with Wnt ligand receptors, Fzd and LRP5/6 at the plasma membrane, which then recruits other components of the destruction complex leading to inactivation of β-catenin degradation. Although this model is widespread, direct evidence for this process is lacking. In this study, we tagged mEGFP to C-terminus of *dishevlled2* gene using CRISPR/Cas9 induced homologous recombination and observed its dynamics directly at the single molecule level with Total Internal Reflection Fluorescence (TIRF) microscopy. We focused on two questions: 1) What is the native size and the dynamic features of membrane-associated Dvl complexes during Wnt pathway activation? 2) What controls the behavior of these complexes? We found that membrane bound Dvl2 is predominantly monomer in the absent of Wnt (mean size 1.10). Wnt3a stimulation leads to an increase in the total concentration of membrane-bound Dvl2 from 0.08/μm^2^ to 0.34/μm^2^. Wnt3a also leads to increased oligomerization which raises the weighted averaged mean size of Dvl2 complexes to 1.4; with 65% of Dvl still as monomers. The driving force for Dvl2 oligomerization is the increased concentration of Dvl2 at the membrane caused by increased affinity of Dvl2 for Fzd, the Dvl2 and Fzd binding is independent of LRP5/6. The oligomerized Dvl2 complexes have greatly increased dwell time, 2~3 minutes compared to less than 1 second for monomeric Dvl2. These properties make Dvl a unique scaffold dynamically changing its state of assembly and stability at the membrane in response to Wnt ligands.

**Significance Statement:** Canonical Wnt signaling is one of the most widely distributed pathways in metazoan development. Despite intense genetic and biochemical study for over 35 years, the major features of signaling across the plasma membrane are still poorly understood. Dishevelled serves as an essential bridge between the membrane receptors and downstream signaling components. Attempts to reconstruct the pathway and analyze its biochemical features in vitro have been hampered by Dishevelled’s tendency to aggregate in vitro and to form large aggregates of dubious significance in vivo. To obtain a molecular understanding of the role of Dvl in Wnt signaling, while circumventing these aggregation problems we have expressed a fluorescent tagged Dishevelled in cells at their physiological concentration and quantified the size distribution of Dishevelled before and after Wnt treatment. We found that limited oligomerization in response to the Wnt ligand is very dynamic and provides a key step of signal transduction.

## Introduction

The canonical Wnt signaling pathway regulates the level of β-catenin, which serves as a transcriptional co-activator. Wnt pathway is a central and conserved developmental pathway in all metazoans. Some of the genes of this pathway originated in Choanozoans but the core pathway is clearly already established in sponges, tens of million years before the Cambrian explosion. In animals the Wnt pathway has a conserved role in: anterior-posterior axis formation in embryos, in organogenesis, and in tissue homeostasis (1–3). Dysregulation of the Wnt pathway plays a role in various human diseases including skeletal, cardiac vascular diseases (4), and cancer (5). Animals generally have multiple Wnt ligands, e.g. 19 Wnt ligands, 10 Fzd and one LRP5, one LRP6 receptor proteins in mammals. In the OFF state of the canonical Wnt pathway, β-catenin levels are low due to active proteolysis mediated by a conserved set of proteins associated in a putative destruction complex (6). Unlike protein complexes such as the proteasome or the anaphase promoting complex, the stoichiometry of the β-catenin destruction complex has never been established, nor has it been purified or crystalized as such. It might very well be unstable or structurally flexible. In the absence of a Wnt signal, the scaffold proteins APC and Axin bind β-catenin and recruit the kinases CK1α and GSK3. A priming phosphorylation on β-catenin mediated by CK1α leads to further phosphorylation by GSK3. The phosphorylated β-catenin is then recognized and ubiquitinated by the E3 ligase β-Trcp, and targeted by the proteasome for degradation. Wnt ligand binding to the Fzd and LRP5/6 receptors at the plasma membrane somehow interferes with this process and therefore causes a buildup of β-catenin protein (the rate of protein synthesis is unaffected by Wnt pathway activation) (7). The Wnt signal for inhibiting degradation is thought to proceed by interference with the activity of CK1α and mainly GSK3. Several molecular mechanisms for the Wnt mediated repression of GSK3 activity have been proposed. These models include that Wnt-induced LRP5/6 phosphorylation acts as a pseudo substrate and directly inhibits GSK3 activity (8), or that Wnt-induced Axin1 dephosphorylation and a conformational change leads to dissociation of β-catenin and Axin1 (9), or that Wnt-induced receptor endocytosis drives GSK3 into Multi-Vesicular bodies, sparing the cytosolic β-catenin from phosphorylation (10).

Apart from APC and Axin, a third scaffold, Dvl (Dishevelled) plays an essential but cryptic role for Wnt pathway activation; its loss completely abolishes the activation of the pathway and leads to constitutive β-catenin degradation regardless of Wnt stimulation (11). But the means by which Dvl carries out its function is still not defined. Unlike APC and Axin, which primarily work as scaffolds allowing for the assembly of the putative destruction complex, Dvl’s only apparent role in canonical Wnt pathway is inhibiting the destruction complex. One popular model is that Dvl bridges the destruction complex and the receptors, communicating the effect of Wnt binding perhaps by interacting with both Fzd and Axin1. In this view Dvl performs its task by binding to intracellular domains of Fzd and forms a receptor complex bringing together Fzd and the co-receptor LRP5/6 in what is known as the signalosome (11). Dvl may stay as a stable complex or shuffle on and off the membrane (8). Dvl may also guide Axin1 and other components to the plasma membrane which may then disrupt the destruction complex and repress its β-catenin degradation activity. Much evidence supporting this model has come from biochemical and imaging experiments with overexpressed Wnt pathway components. While approaches based on protein overexpression can be informative, they would have potential limitations if concentration is important to the properties of the system. This state of uncertainty ultimately may only be clarified by direct visualization of dynamics of pathway components at physiological levels.

In studies of pathway dynamics, the importance of expressing proteins in cells at their physiological levels cannot be overemphasized. Overexpression, which is widely used, comes with caveats of potential artifacts. The Wnt pathway scaffold proteins have dimerization (12, 13) or oligomerization (14, 15) domains, which carry with them the potential of forming large non-physiological structures when expressed at high concentration (16–18). Whether these structures exist in the physiological context remains a matter of debate. Specifically, the DIX domain of Dvl exhibits head-to-tail oligomerization, and overexpressed Dvl form so called “puncta” (15, 19–22), which are protein structures as big as 1~2 μm in some cases (over a million Dvl molecules if composed with only Dvl). A physiological function for these “puncta” of Dvl has been proposed as a scaffold to repress the destruction complex by some groups (15, 21) but challenged by others (23). With the help of modern CRISPR approaches for endogenous tagging of proteins, we are now able to ask questions of the form of Dvl when not overexpressed. With TIRF (Total Intensity Reflection Fluorescence) Microscopy, it is possible to measure dynamic processes at the sub-second time scale with a spatial resolution that allows the resolution of individual molecular sized complexes. Using such approaches, we have addressed anew the size and dynamic features of membrane-bound Dvl complexes during Wnt pathway activation and analyzed the control of Dvl binding at the plasma membrane.

## Results

### Most Dvl2 is evenly distributed in the cytosol regardless of Wnt3a treatment

To understand the distribution of Dvl, we fused mEGFP to the C-terminus of *dishevelled2* gene in HEK293T cells using CRISPR/Cas9 mediated genome-editing. HEK293T cells express all three Dvl isoforms; Dvl2 constitutes more than 80% of the total pool (24), indeed Dvl2 deletion results the most severe phenotype in mouse models (25). In knock-in cells the concentration of Dvl2-mEGFP is less than 8% different to the concentration of the Dvl2 in WT HEK293T cells, and the response of β-catenin kinetics to Wnt3a is nearly identical in the two cell lines (Figure 1A). We imaged Dvl2-mEGFP knock-in cells for 10 hours by confocal microscopy after the cells were activated by Wnt3a. Dvl2-mEGFP was evenly distributed in the cytosol and rarely found in the nucleus or on the membrane. (Figure 1B, movie S1). In a few cells, some Dvl2 is localized to a structure that might correspond to the centrosome (26) or lysosome (27). The lack of any discernable plasma membrane localization is likely to be due to the high background of cytosolic Dvl2 and the limited sensitivity of the CCD chip in the confocal microscope. Indeed when we overexpressed snap tagged SNAP-Fzd5 in the same cells (Figure S1), we found some Dvl2 complexes clearly co-localized with SNAP-Fzd5 at the plasma membrane. Presumably the number of Dvl2 complexes on the membrane before or after Wnt3a treatment is very low in HEK293T cells either due to low density of membrane Fzd or weak interaction between Dvl and Fzd. Given such strong cytoplasmic background, it is likely that such a small number of complexes would be very difficult to be observe.

**Figure 1.**
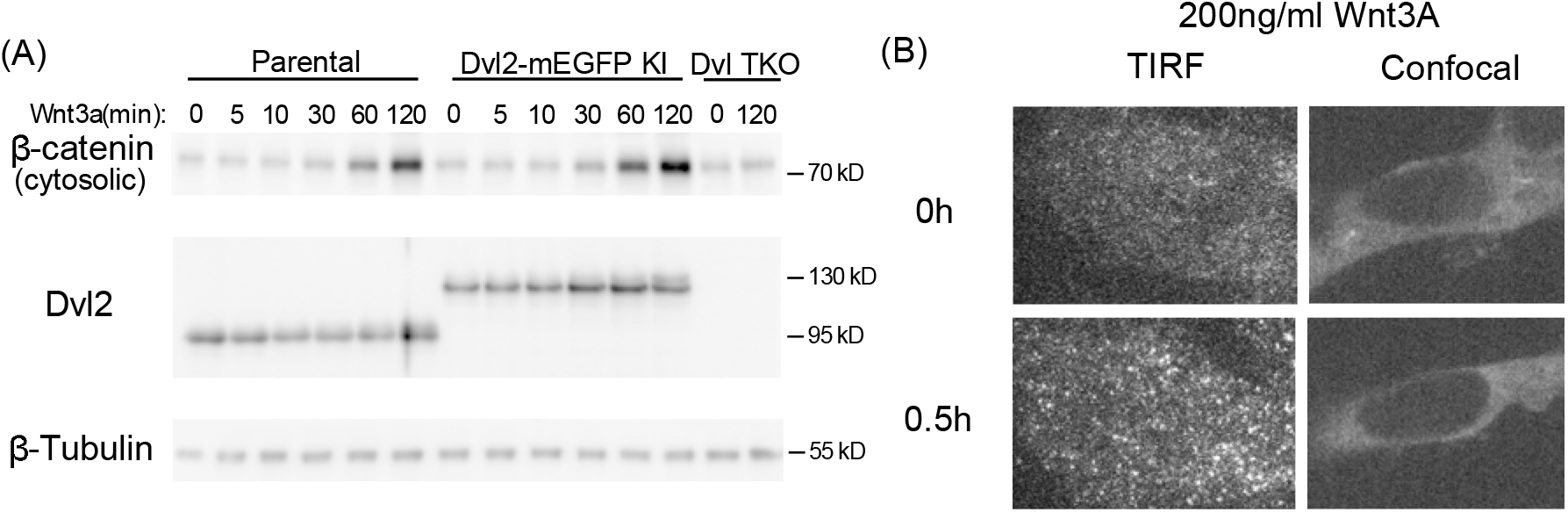
Dvl2 forms complexes on the cell membrane in response to Wnt3a treatment. (A) β-catenin level increase and Dvl2 concentrations are similar in HEK293T cells and Dvl2-mEGFP knock-in cells in response to Wnt3a treatment for 2h, but not in Dvl-TKO cells. (B) Dvl2-mEGFP under TIRF and confocal microscope.

### TIRF visualization and quantification of Dvl2 oligomer states on the plasma membrane before and after Wnt addition

To overcome the limitations of confocal microscopy, we imaged the labeled Dvl2-KI cells using Total Internal Reflection Fluorescence (TIRF) microscopy. TIRF microscopy illuminates a depth about 150 nm above the coverglass, which greatly reduces the fluorescence glow from the cytosol. Dvl2-KI cells were imaged once every 30 seconds for 75min; Wnt3a (200ng/ml) was added to the cell culture media at 15min and images were processed (28) (an example of single molecule detection shown in Fig. S2) to extract the position and intensity of the fluorescent spots and to link fluorescent spots in neighboring frames. To determine the number of Dvl2-mEGFP in each complex, we used two methods to quantitate single mEGFP intensity. The first by analyzing the intensity distribution of all the light spots in one image, and the second by analyzing precipitous jumps of intensity in all the single molecule traces, either due to photobleaching or to association/dissociation of Dvl2-mEGFP subunits. The absolute value of all the light spots (Fig. 2C) or intensity jumps (Fig. 2B, 2D) were compiled into histograms and fitted with a mixture Gaussian function to establish the single mEGFP intensity value. The ratio of mEGFP intensity value achieved from the two methods is 1.09±0.16 (6 control data sets). The first method does not require high quality long single molecule traces, so we use it principally in later analyses to quantify the mEGFP intensity. Additionally, measurement of another membrane protein mEGFP-GlyRa1 yielded similar single mEGFP intensity value (Fig. S3).

**Figure 2.**
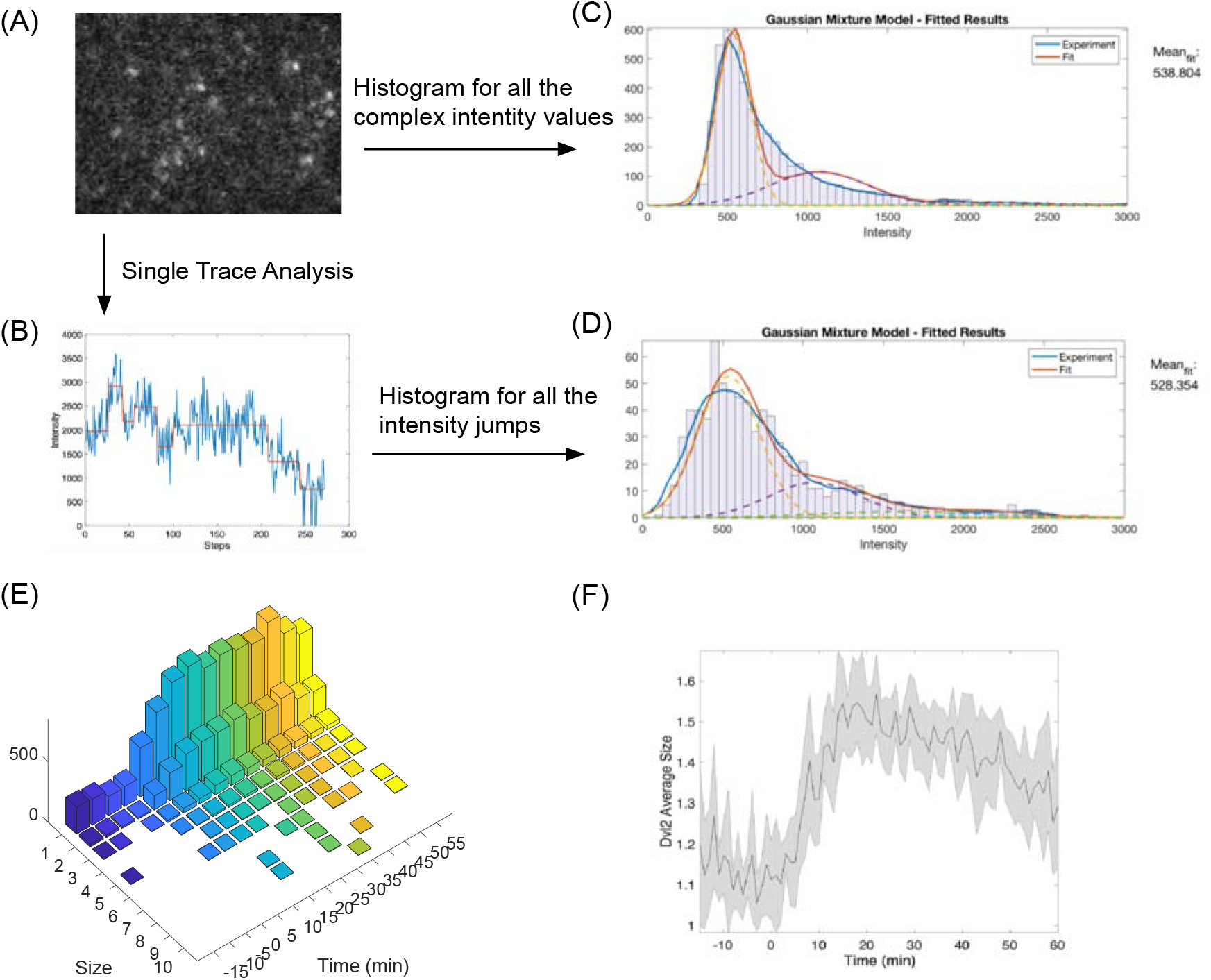
Quantification of the size and dynamics of membrane Dvl during Wnt pathway activation. (A) single molecule images are analyzed and single molecule traces (B) are used to find steps where mEGFP are added or removed/photobleached. Either the intensity of all the complexes in the image (C) or all the steps in the single molecule traces (D) are analyzed to extract single mEGFP intensity value. (E) The time course of Dvl2-mEGFP size distribution is then quantified with 200 ng/ml Wnt3a added at time 0. (F) Average size of Dvl2-mEGFP complexes only increase from 1.1 to 1.4.

In the absence of Wnt, we found that 89.2±3.9% of membrane Dvl2 complexes contain only a single Dvl2, with 10.7±3.7% containing two molecules of Dvl2, virtually none of the spots have more than two Dvl2 molecules. Increased Dvl2 molecules localize to the cell membrane within 5 minutes of Wnt3a treatment and reach a maximum at 15min, gradually declining to a level that is higher than before Wnt3a treatment (Fig. 2E). This time frame is very similar to that for the decrease of fully (T41/S37/S33) phosphorylated β-catenin observed in bulk assays; this is still before there is a measurable increase in β-catenin concentration (30 min) (7). Even following Wnt treatment, the complexes that contain monomeric Dvl2 are still the majority population, comprising 65.4±2.3% of the total complexes. There is, however, a significant shift to larger complexes. By 15 min 26.8±2.3% of the complexes contain two Dvl molecules and the rest (18%) more than two. As a result, the weighted average size of Dvl oligomers increases from 1.1 to 1.4 (Fig. 2F).

To estimate the number of Dvl2 bound to the plasma membrane, we start with the cellular Dvl2 concentration in a single HEK293T cell, which we estimate to be 140 nM (Fig S4). With a cell volume 1.94 pL measured by ORFLO^®^, on average there should be a total of 1.6×10^5^ molecules of Dvl2 per cell. Our microscopic measurements allow us to estimate the area density of Dvl2 to be ~0.3/um^2^ (Fig. 3B) after Wnt3A treatment. Assuming the cell to be spherical, its total surface area would be about 7.5×10^2^ μm^2^ (note that a model of a flattened cell as an oblate ellipsoid of revolution with an axial ratio of 4 and the same volume of the cell would have surface area about twice as large, 1.6×10^3^ μm^2^). On the membrane therefore there should be between 2.3×10^2^ molecules of Dvl for the spherical cell model to 4.8×10^2^ molecules of Dvl for the oblate ellipsoid of revolution cell model. Therefore, less than 0.3% of the cellular Dvl2 is on the membrane, even at peak Dvl2 recruitment. This small fraction of Dvl at the membrane together with high cytosolic Dvl2 background explains why membrane Dvl was invisible by confocal microscopy but why we could detect it by TIRF microscopy.

**Figure 3:**
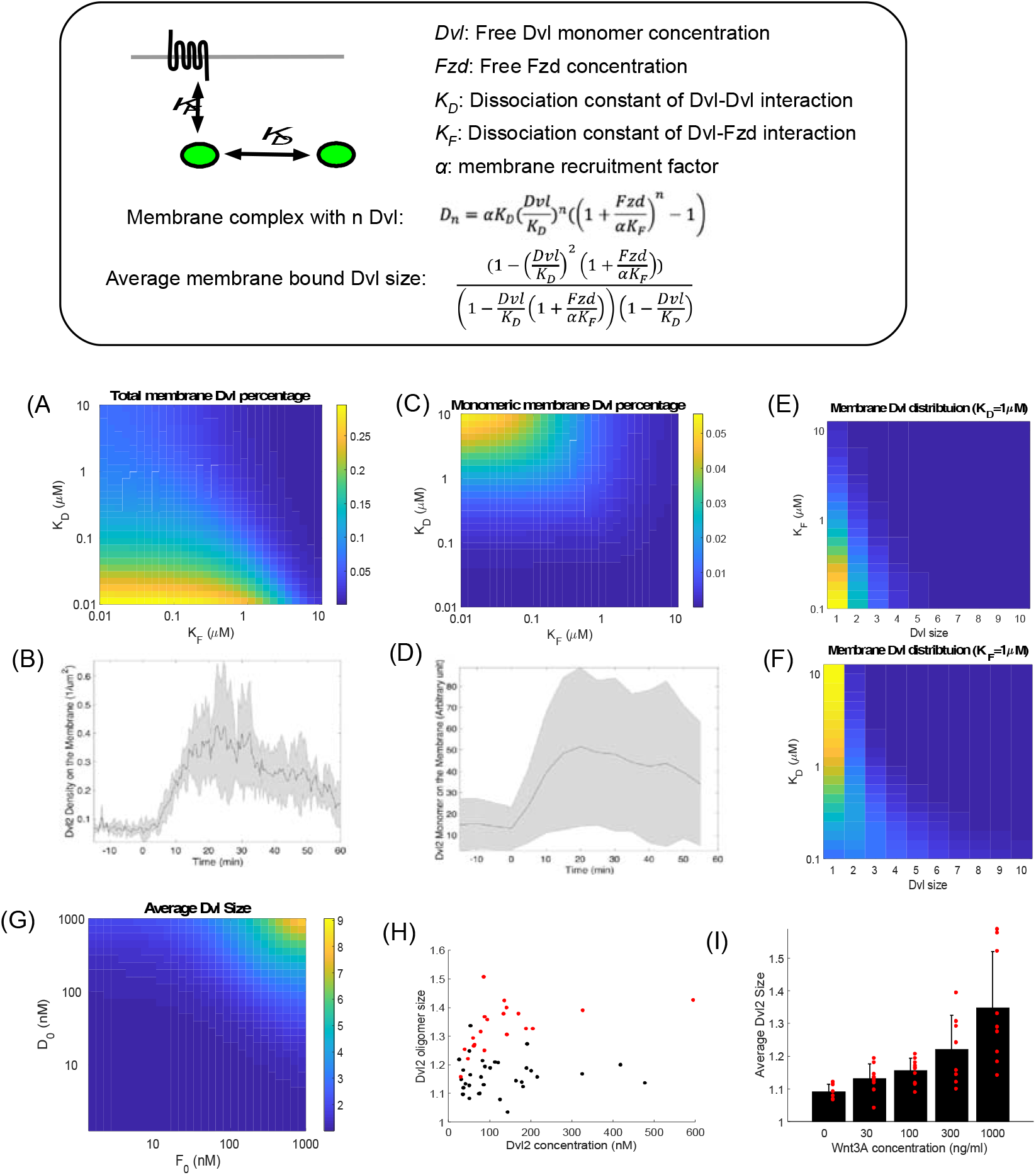
Dvl membrane recruitment is driven by Dvl-Fzd interaction. (A) Predicted percentage of membrane bound Dvl while varying *K*_*D*_ and *K*_*F*_. (B) Experiment data for Dvl2 membrane density with Wnt3a treatment. (C) Predicted monomeric Dvl on the cell membrane when varying *K*_*D*_ and *K*_*F*_. (D) experiment data for membrane monomeric Dvl2 with Wnt3a treatment. (E) Predicted Dvl size distribution with *K*_*D*_ fixed while varying *K*_*F*_. (F) Predicted Dvl size distribution with *K*_*F*_ fixed while varying *K*_*D*_. (G) Predicted average size of Dvl complexes while varying Dvl and Fzd concentration. (H) average size of Dvl complexes while changing total Dvl2 concentration. Each dot corresponds to a single cell. Black dots are cells without Wnt3a treatment while red dots are cells with Wnt3a treatment. (I) average size of Dvl complexes while changing Wnt3a concentration.

### Mass action driven reactions reveal increased Dvl-Fzd interactions after Wnt treatment

The change in oligomerization of Dvl on the membrane in response to Wnt3a is likely due to homotropic interactions between Dvl molecules and heterotropic interactions between Dvl and Fzd. Since TIRF microscopy can quantitatively measure the size and distribution of Dvl membrane complexes, we have tried to explain the Dvl changes with the simplest possible model to understand the perturbation by the Wnt ligand. Our basic model contains only Fzd and Dvl, where Fzd is the membrane receptor that binds Dvl. Dvl could form head-to-tail linear oligomers mediated through its DIX domain (29). Thus, we consider two basic binding reactions in the model, Dvl-Dvl binding and Dvl-Fzd binding. We define the dissociation constant between Dvl molecules to be *K*_*D*_, and the heterotypic dissociation constant between Dvl and Fzd to be *K*_*F*_. Once on the membrane, proteins might have different binding affinities due to changed local concentration and membrane induced constraint on protein structure. We do not explicitly calculate the membrane local concentration, instead we assume any Dvl complexes that bound to Fzd are on the membrane. We signify the membrane effects on both association constants by adding a factor α (<1) to indicate the increased binding affinity of Dvl with other molecules once bound to Fzd. We then have the following equilibrium equations:

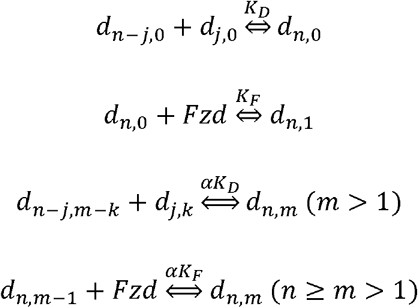

Here *d*_*n,m*_ represents the concentration of *one specific configuration* of Dvl complexes that contains *n* Dvl molecules and *m* Fzd molecules. Note that *d*_*n,0*_ contains 0 Fzd thus represents cytosolic Dvl complexes. The first equation describes Dvl complex formation in the cytosol. The second equation shows the recruitment of cytosolic Dvl complexes to the membrane by Fzd. The third equation describes the combining and splitting of membrane bound Dvl complexes, and the fourth equation denotes the binding and unbinding of membrane bound Dvl complexes to Fzd. α in the last two equations represents general membrane effects. At equilibrium, we can solve for the concentration of all forms of Dvl complexes (supplement text S1). Note that there are only 5 parameters in the model, and we can fix Dvl concentration with our experimental measurements. This allows us to explore the behavior of the model in the full parameter space.

In single molecule imaging experiments, we found that Dvl is increasingly localized to the membrane during Wnt pathway activation. This could be explained by an increase in Dvl protein concentration, or an increase in its binding affinity. When we measured the concentration of Dvl by Western blot in response to Wnt; there was <20% change in the 4-hour period after Wnt treatment (Fig. S5). During this time there was a 30~40% decrease of Fzd during the first hour of Wnt activation (as measured by the level of overexpressed surface mEGFP-Fzd1 with TIRF microscopy, Fig. S6). Given little change in total cellular Dvl, the remaining possible explanation for increased Dvl at the membrane must due to a decrease of either *K*_*D*_ or *K*_*F*_. To explore the model further we fix α, *Dvl*_*0*_ and *Fzd*_*0*_ and ask how the changes in *K*_*D*_ and *K*_*F*_ affect the amount of Dvl at the membrane.

The parameter α changes the apparent concentration on the membrane and in turn determines the degree by which the membrane environment affects both Dvl-Dvl binding and Dvl-Fzd binding. We tried α=0.1~0.01 and found that α does not affect the general shape of the membrane Dvl size distribution. For further analysis of the oligomerization of Dvl we choose an arbitrary value for α, which we fixed at 0.01. Dvl concentration is 140 nM, as measuredexperimentally. We assumed the Fzd concentration to be 10 nM; higher or lower concentrations does not qualitatively change the following conclusions. Not surprisingly the model predicts that a decrease (increase the binding affinity) of either *K*_*D*_ or *K*_*F*_ can result in increased membrane bound Dvl (Fig. 3A), as we observed in our experiments (Fig. 3B). However, by computing the average Dvl complex size from the model (Fig. S7), we find that the average size increases to more than 2 when *K*_*D*_ is in the sub μM range, indicating weak Dvl-Dvl interactions in the cell. Furthermore, we found that to explain the large increase in monomeric membrane Dvl there must be a decrease (increased binding affinity) in *K*_*F*_ but not in *K*_*D*_ (Fig. 3C, 3D). This systematic search of parameter space strongly suggests that Wnt treatment acts primarily by increasing the binding affinity between Fzd and Dvl (a decrease in *K*_F_).

The size distribution of Dvl contains rich information about the exact molecular interactions. We find that *K*_*D*_ controls the shape of the distribution while *K*_*F*_ controls the height of the distribution. Decreased *K*_*D*_ favors large Dvl oligomers which flattens the distribution, while increased *K*_*D*_ skews the distribution to the monomeric Dvl side. On the other hand, a decreased *K*_*F*_ would increase Dvl-Fzd binding which in turn increases the total membrane-bound Dvl, resulting in the distribution shifting upward, while a larger *K*_*F*_ would move the distribution downward. By comparing the experimental distributions (Fig. 2E) to theoretical ones (Fig. 3E, 3F), we conclude that the shift-up of the experimental distribution after Wnt3a treatment must be due primarily to decreased *K*_*F*_(stronger Dvl-Fzd binding). A more complete set of simulations covering larger parameter range can be found in Figure S8. Based on the ratio of membrane Dvl to total Dvl, and the shape of the distribution, we estimate that in the absence of Wnt, both *K*_*D*_ and *K*_*F*_ are weak, in the μM range.

The model also predicts that increased concentration of either Dvl or Fzd should result in an increased average Dvl oligomer size (Fig. 3G). Presumably an increase in Dvl concentration will increase the oligomerization of Dvl through its DIX domain, therefore increasing the oligomerization of Dvl in the whole cell, including the membrane. An increased Fzd concentration, on the other hand, increases the membrane local density of Dvl which will be the driving force to form larger oligomers. Both predictions can be verified experimentally. To vary Dvl concentration in the cell, we over-expressed Dvl2-mEGFP in Dvl-TKO cells, which has all three Dvl knocked out (30), and measured cytosolic concentrations with wide-field fluorescence microscopy and Dvl2 size on the membrane with TIRF microscopy. We found a weak but positive correlation between increased Dvl2 size and an increased Dvl2 total concentration. Dvl2 size further increases in response to Wnt3a (Fig. 3H). We use an indirect way to change apparent Fzd concentration by adding different concentration of Wnt. As predicted by the model, Wnt3a should induce a population of Fzd with higher binding affinity to Dvl. Indeed, increased Wnt3a concentration produces a higher average Dvl oligomer size on the plasma membrane (Fig. 3I).

### Structural requirements for Dvl plasma membrane recruitment

In principle Dvl could be recruited either by direct interactions with Fzd or through other intermediate proteins. Dvl is known to have three globular domains: DIX, PDZ and DEP. It has been shown recently that the DIX and DEP domains in Dvl are important for canonical Wnt pathway function (31). The PDZ domain has also been reported to bind directly to the C-terminus and inter-helix loop region of Fzd (32). To test which domains affect the membrane dynamics of Dvl, we individually removed the DIX, PDZ or DEP domains from the Dvl2-mEGFP construct and assayed the effects on Wnt activation, using a transcriptional TopFlash assay (Fig. 4A-B). When we overexpressed ΔDEP-Dvl2 in Dvl-TKO cells, it failed to rescue the Wnt pathway activity. The ΔDIX mutant showed a very low degree of complementation. Notably expression of ΔPDZ-Dvl2 strongly complemented the knockout, but not to the original level. There were corresponding effects of expressing these constructs on Dvl recruitment. In TIRF measurements for the above cell lines, we found that cells expressing ΔDEP-Dvl2 failed to recruit additional Dvl to the membrane on Wnt3a addition and the Dvl on the membrane did not increase its degree of oligomerization. We also found the membrane density of ΔDEP-Dvl2 to be lower than WT-Dvl2. By contrast the ΔDIX-Dvl2 complemented the membrane Dvl density compared to WT-Dvl2 but did not increase the level of binding or oligomerization on Wnt addition. Dvl with PDZ domain deletion behaves similarly to WT-Dvl both in the level of binding and the increased membrane localization and oligomerization in response to Wnt (Fig. 4E-F). These properties of the Dvl2 mutants attest to the importance of Dvl membrane recruitment in Wnt pathway activation and the central importance of the DIX and DEP domains.

**Figure 4.**
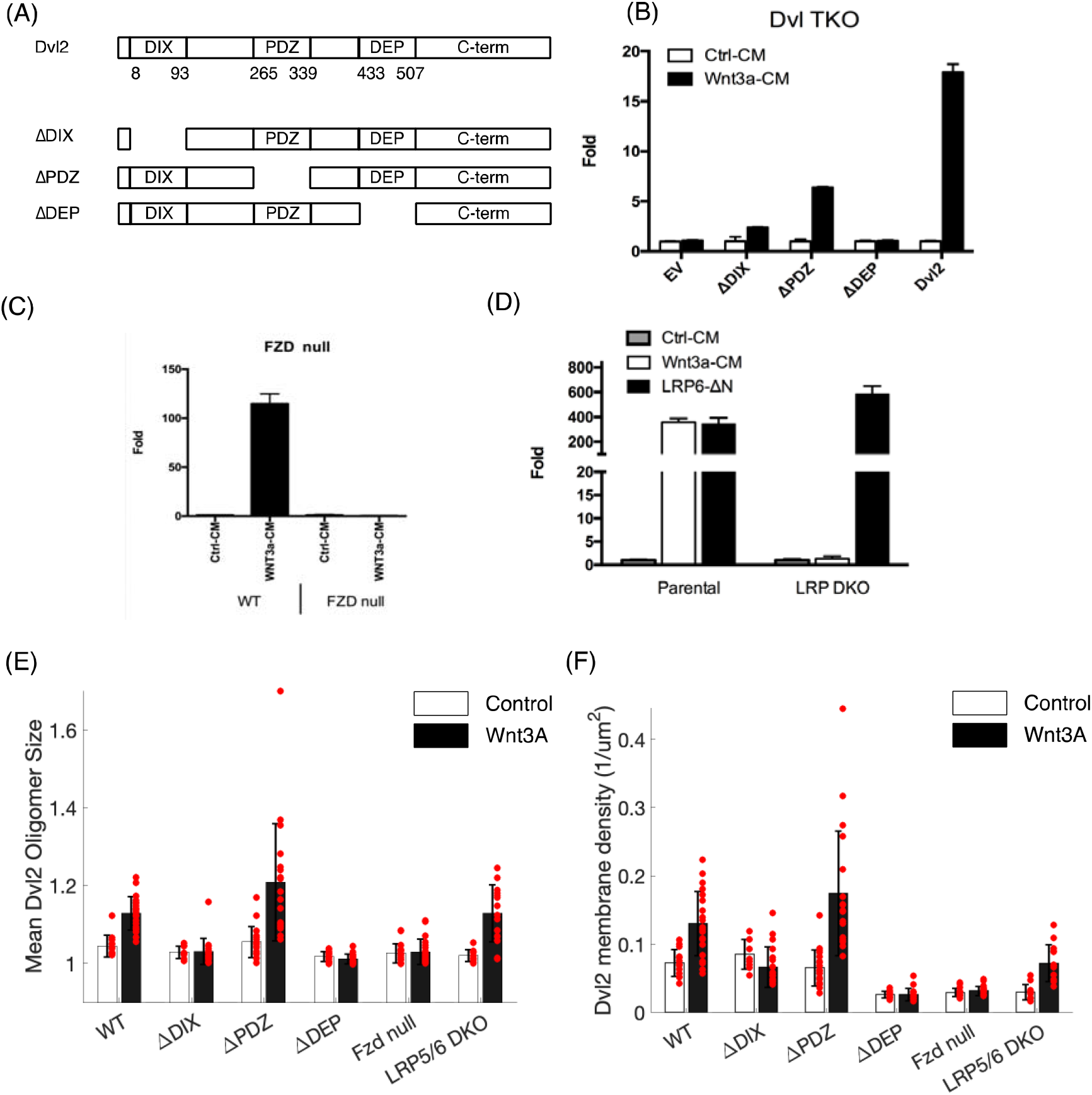
Dependence of Dvl membrane localization to the Dvl protein domains and receptors. (A) Dvl2 mutations used in the analysis. (B) Top-flash response of different Dvl2 mutants. (C) Top-flash assay of Fzd-TKO cell. (D) Top-flash assay of LRP5/6 KO cell. (E) Mean Dvl2 number per complex under different conditions. Each dot represents a cell. The error bars represent standard deviation. (F) Dvl2 membrane density under different conditions. The error bars represent standard deviation.

Similar experiments allowed us to test whether the membrane recruitment of Dvl depends on Fzd or LRP5/6. We made Dvl2-mEGFP knock-in cell lines in an Fzd-null (all Fzd knockout) cell line; and similarly in an LRP-DKO (LRP5/6 double knockout) cell line. In cells lacking Fzd, we found no Dvl membrane recruitment upon Wnt treatment, in agreement with our prediction that Fzd-Dvl interaction increases after Wnt treatment. By contrast in LRP5/6 double knock-out cells (Chen & He, Dev Cell, in press), Dvl2 still increases its localization to the plasma membrane after Wnt treatment, although it fails to activate downstream genes (Fig. 4). Furthermore, while canonical Wnt pathway inhibitor, DKK1 blocks Wnt induced β-catenin accumulation through direct binding to the Wnt3a binding region of LRP5/6, cells still show increased Dvl2 membrane localization and oligomerization (Fig S9). Taken together, these experiments show that Dvl membrane recruitment is Fzd dependent but LRP5/6 independent. Dvl is known to be a general mediator of both canonical and non-canonical pathways. Therefore, it is reasonable to expect that Dvl response is dependent on Fzd but not on the pathway specific co-receptors, such as LRP5/6 and Ror1/2 (33).

### Measurement of the dwell time of Dvl2 on the membrane and its potential significance

Although steady state descriptions are useful, many complex processes are also governed by the dynamical processes. This is especially true if such processes involve chemical reactions, such as phosphorylation or ubiquitination, where accumulation of the modification is tied to the time an *individual* molecule remains on the site rather than the average occupancy of that site (34). For this reason, we extended our study of the steady state distribution of Dvl on the membrane to examine the dwell time of *individual* Dvl complexes. To analyze the dwell time of individual membrane bound Dvl2 oligomers, we made 20Hz videos with TIRF microscopy and tracked the protein complexes with uTrack. These video images show the diffusional movement of the molecule in the plane of the membrane where the persistence of the signal is registered by the number of frames that contains a specific protein complex, thus providing the measure of dwell time of Dvl in the membrane complex. One important complication is that in addition to dissociation of Dvl from the membrane, photobleaching can also terminate the signal, especially when images are taken at high imaging rate with a high intensity laser. To correct for the effects of photobleaching, we used a membrane bound pentameric channel protein GlyRa1-mEGFP as the control to estimate photobleaching time scale (Fig. S10), where the probability distribution of photobleaching can be approximated as geometric distribution. We estimated the photobleach rate constant to be 1.17±0.13/sec, which corresponds to mean half-life of 0.59 second (11.8 frames) for mEGFP.

Two potential mechanisms could change the dwell time of Dvl complexes. Dissociation from the membrane will terminate the presence of Dvl complexes in the series of video frames thus shortening the trace length. On the other hand, merging with other membrane bound or new Dvl molecules from the cytosol would extend the trace length (Fig. 5A). Based on these basic processes, we showed by simulation that the dwell time distribution of photobleaching can be modeled by a geometric distribution which decays exponentially with increased dwell time. The distribution would still be geometric after adding the dissociation term, but the experimental traces would have considerably shorter dwell times. Adding merging and splitting of Dvl molecules to the model lengthens the apparent dwell time. The reasons are twofold, first merging Dvl complexes introduces fresh fluorophores to the membrane complexes that counteracts photobleaching; second the resulting larger complexes have a reduced rate of dissociation. (Fig. 5B).

**Figure 5.**
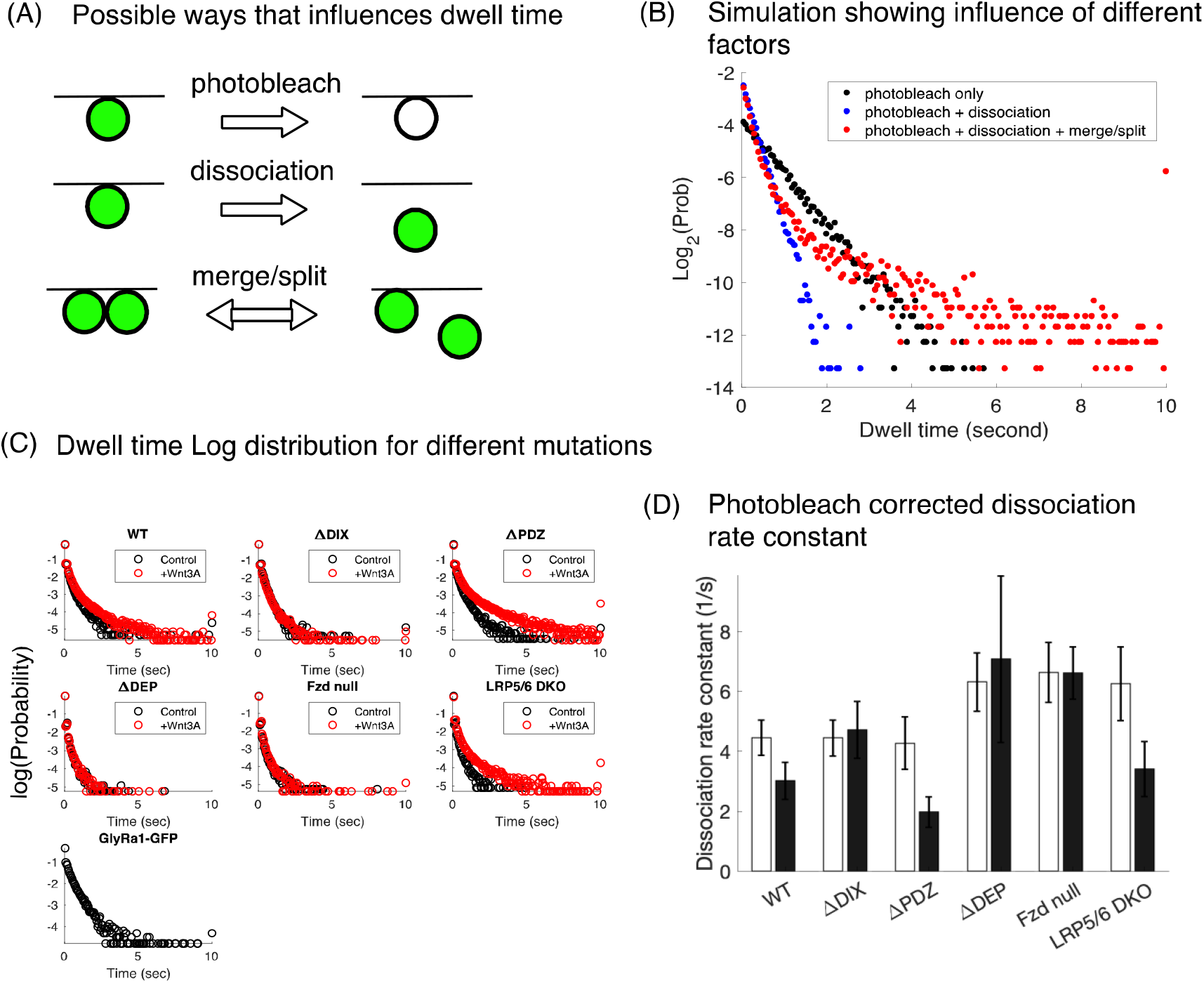
Dwell time analysis of membrane bound Dvl. (A) Possible mechanisms that changes the single molecule dwell time measurement. (B) Simulations to show how different mechanisms affecting dwell time distribution. (C) Pooled dwell time Log distribution for different mutant cell lines. (D) Dissociation rates derived from the upper part of dwell time distribution. The error bars represent standard deviation.

With this framework, we can begin to estimate the dissociation rate for all the mutants and Fzd KO, LRP5/6 DKO cells. The pooled dwell time distributions of different mutants are shown in Figure 5C. Wnt3a clearly shifts the distribution to form a long tail for WT-Dvl2, Dvl2-ΔPDZ in Dvl TKO cells, and in LRP5/6 DKO cells, indicating active Dvl-Dvl association and dissociation events, and consistent with the requirement of the DIX domain for Dvl-Dvl interaction and the DEP domain for Dvl-Fzd interaction. We used the steep decline region of the dwell time distribution to approximate the dissociation rate constant (Supplement Text S2), which is shown in Figure 5D. The fitted dissociation coefficients are in the range of 2~10/sec. The frame rate of the TIRF microscope (20/sec) limits the precision of our estimate of the dissociation constants for these fast dissociation events. Nevertheless, if we assume the association rate constant to be 5×10^5^/M/sec for proteins the size of Dvl, the dissociation constant will be in the range of 4~20 μM, which is very weak at a Dvl2 concentration of 140 nM. This estimate of the dissociation constant confirms our estimate based on the modeling of Dvl-Fzd association. Such weak binding explains the small number of Dvl on the plasma membrane (less than 0.3% of the cytosolic concentration) despite relatively high Dvl concentration in cytosol. In the presence of Wnt3a the average dissociation coefficient for WT-Dvl2 changes from 6/sec to 4/sec, indicating increased interaction between Dvl and the membrane, presumably through Fzd. This further confirms our analysis of the Dvl size distribution and membrane localization. The change of dissociation coefficient is less than two-fold because the number reflects the pair-wise interaction change between Dvl and Fzd for a mixed population of Fzd, both Wnt bound and Wnt free. As expected, we found Dvl without the DEP domain increases the dissociation coefficient from ~6/sec to ~10/sec, as do Dvl2 in Fzd KO cells, indicating decreased Dvl membrane binding. These mutants also show no change in dissociation constant with Wnt3a treatment. Surprisingly LRP5/6 knock-out cells show decreased Dvl membrane binding but still respond to Wnt3a. This suggests that in the absence of LRP5/6 the Wnt3a interaction with Fzd is enough to affect the binding affinity of Fzd with Dvl. Dvl-ΔDIX mutant shows similar basal membrane binding with wildtype Dvl but does not respond to Wnt3a.

Due to photobleaching, the averaged dissociation rate estimate in Fig. 5D reflects the time scale of the population with the shortest dwell time. To estimate the dwell time of the relatively few large Dvl complexes that bind more stably to the membrane, we measured their fluorescence recovery after photobleaching. We imaged the recovery process by using 5s time intervals for 10min after photobleaching, where the fast time scale ~5/s would not show up. This experiment showed that the recovery time is indistinguishable before (125±65s) and after (156±96s) Wnt3a treatment. Arguably this population of large stable complexes would be stable before and after Wnt addition. The dwell time will become of more than academic interest, if relevant chemical reactions occur at similar time scale. Dvl is well known to be phosphorylated at multiple sites (35), probably at the plasma membrane, although the exact function of these modifications is unclear. Based on the BRENDA database, which catalogs enzyme kinetic information, the turnover number of general Ser/Thr kinase like CK1 is only 0.057~0.19/s, slower than the fast time scale of Dvl-Fzd interaction (4~6/s) discussed above, but faster than the slow time scale which corresponding to the average Dvl oligomer binding (0.0083~0.0056/s). If Dvl phosphorylation happens on the membrane, the transient binding and low membrane stoichiometry of Dvl means there would be a very low probability of Dvl phosphorylation. However, after Wnt pathway activation and the formation of Dvl oligomers, there could be appreciable phosphorylation, as the average dwell time increases from less than a second to a few minutes. A process dependent on dwell time rather than occupancy would ensure a very low background of Dvl phosphorylation in the absence of Wnt. While the effect of Wnt may seem modest, it is sufficient to switch a significant fraction of Dvl complexes from the unphosphorylated state to an active phosphorylated state.

### How oligomerization affects Dvl Dynamics and activates the Wnt response

The oligomerization of Dvl complexes appears to be a *necessary* feature of canonical Wnt pathway activation. Can we say something about whether association of Dvl subunits itself is *sufficient* to activate the pathway? To answer this question, we generated a series of artificial Dvl oligomers by replacing the DIX domain of Dvl2 with a set of constructs containing unrelated oligomerization motifs, artificially designed to generate specific oligomer states (36). The set of Dvl oligomeric constructs are named Dimer, Trimer, Tetramer, and Hexamer which correspondingly contain 2, 3, 4, and 6 Dvl molecules in stable complexes (Fig. 6A). The size of the various stable oligomeric constructs was further confirmed by sucrose gradient sedimentation (Fig. S11). Each of these modified forms of Dvl was separately expressed in Dvl-TKO cells and the activation of the Wnt pathway was measured by the TopFlash assay. Wnt pathway activation peaks at the Dvl trimer and drops as the Dvl2 size further increases (Fig. 6B). This result suggests that there is an optimum in the stability of complexes, which taken literally means that activity depends on both efficient binding and efficient release.

**Figure 6.**
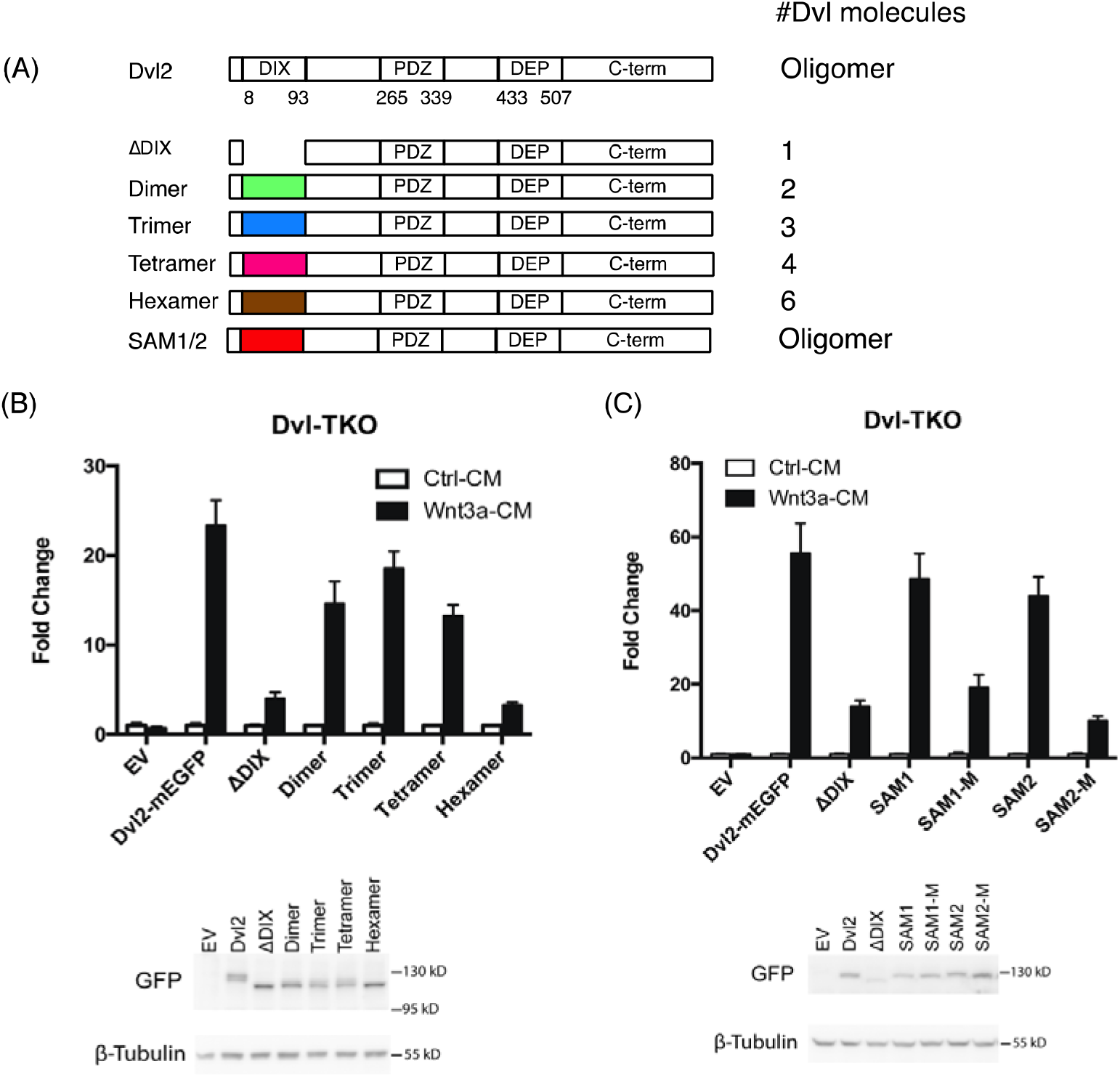
Both Dvl membrane recruitment and Dvl subunit dynamics are important for Wnt pathway activation. (A) DIX domain of Dvl2 are replaced with designed domains that have specific number of subunits. (B) All the mutants have lower activity compared to wildtype Dvl2 but the activity peaks at Dvl2 trimer. (C) DIX domain of Dvl2 is replaced with SAM domain in two species of plants. The SAM domains that can oligomerize successfully recover the activity of Wnt pathway, but the SAM domains with monomer mutations could not.

If the *dynamic* oligomerization of Dvl2 is important for its function, could we replace DIX domain with other domains with similar property and rescue Dvl2 protein function? For this question we used the SAM (Sterile Alpha Motif) domains of LEAFY transcription factor in flowering plants, which also has a tendency to form head-to-tail oligomerize like Dvl DIX domain but do not generate well-defined stable oligomers (37). Interestingly, by replacing DIX domain of Dvl2 with two different SAM domains in the plants (*Arabidopsis thaliana* for SAM1 and *Ginkgo biloba* for SAM2), we show that the Wnt pathway activity can be rescued (Fig. 6C). SAM domain point mutations that disrupt the oligomer forming potential disrupt Wnt pathway activity (SAM1-M, and SAM2-M in Fig. 6C). The previous two experiments suggest that the sole function of the DIX domain on Dvl is to form dynamic oligomers and which activity potentially maximize at trimeric molecules.

### The role of Dvl dynamics in non-canonical Wnt pathways

Dvl is involved in both the canonical Wnt signaling pathway and at least two non-canonical pathways, the Wnt/PCP (Planar Cell Polarity) pathway and Wnt/Ca^2+^ pathway (38). Does Dvl play a role in choosing the downstream responses for these very different circuits and if so, is it related to different dynamics of Dvl oligomers? Although we cannot definitively answer the first question at present, we can address the second question. Wnt3a is a typical canonical Wnt pathway ligand, while Wnt5a is a typical Wnt/PCP pathway ligand. When we examine the response of Dvl2-mEGFP KI cells to the two Wnt ligands, we found that they generate different dynamical responses at the membrane. Although both ligands drive Dvl to the cell membrane and cause an increase in Dvl oligomerization, Wnt5a consistently produces much larger complexes (Fig. S12). Furthermore, when we analyzed the size distribution of Dvl, we found that instead of a near exponential distribution like Wnt3a, Wnt5a treated cells shows a distribution that cannot be fit by a single exponential. The long tail of the Dvl distribution in Wnt5a cells reveals complexes containing 9 to 13 molecules of Dvl. (Fig. S13-S14). The long tail implies some positive feedback that would stabilizes larger Dvl complexes apart from simple membrane recruitment by Fzd. This positive feedback might be caused by increased DIX-DIX interaction or interaction with other proteins or cellular organelles. In cells activated by the Wnt/PCP pathway, Dvl is known to preferentially localize to one side of the cell where it forms extensive patches at the plasma membrane (39). The much larger Dvl complexes that we see generated by Wnt5a may be related to this. To function as the critical signaling component that differentiates the Wnt3a and Wnt5a pathways, Dvl2 would have to show different behaviors with different Wnt ligands. We can now expand the list of distinct behaviors of Dvl from very weak binding of virtually only monomers when there is no Wnt present, to small and dynamic oligomers in Wnt3a related complexes, to larger and more stable complexes for Wnt5a. It remains to be seen if the oligomerization is the only or most important differentiating effect linking different Wnt ligands to downstream pathways.

## Discussion

Our understanding of cellular mechanisms has benefited from biochemistry, carried out in isolated systems approximating the cellular milieu, and from genetics, where the interpretations though indirect, are achieved by definition in a physiological context. When biological processes are uncomplicated and in particular when the products can be isolated as simple chemicals, biochemistry excels. But when the processes are complex and the reaction products cannot be isolated the more indirect genetic approaches have proven to be particularly useful. For many cellular reactions and especially for processes of information transfer in signal transduction where linkages are weak (40), it has not been possible to reconstitute the reactions. Even when there are strong clues from genetics the mechanisms sometimes have often been very difficult to establish in vitro. That describes the state of our understanding of Wnt signaling, one of the most important developmental and oncogenic pathways. The current mechanistic understanding focuses around “destruction complex” and “signalosome”, which are still hypothetical assembly of enzymatic functions, including kinases and ubiquitin ligases, scaffolds that interact with each other in unclear ways and with unknown stoichiometries. Among the most enigmatic among all the components is the essential Dvl scaffold proteins. We have carried out a kinetic study of its behavior in living cells combining CRISPR induced genome modification with single molecule imaging, and analyzed the data through quantitative modeling. This has shed new light on how Dvl may regulate the downstream signaling machinery. These studies point to the importance of characterizing weak dynamic interactions and considering off rates in addition to stoichiometries of binding.

We combined single molecule imaging at the plasma membrane with modeling to study Dvl dynamics during Wnt pathway activation. Our results point out the critical role of Dvl oligomerization at the membrane and the weak binding between Dvl-Dvl and Dvl-Fzd. Given these weak and dynamic interactions it was very important to express labeled proteins in the cell at close to their endogenous levels. We found that the state of Dvl in the cell is defined by its endogenous concentration, which at 140 nM is significantly lower than DIX domain binding affinity to Fzd in the membrane (~μM). We extracted the affinity from a simple model with only Dvl-Dvl and Dvl-Fzd interactions. The disparity of the very low concentration and the weak binding affinity guarantees that the predominant species of Dvl in the cell is in the form of a soluble monomeric pool. The weak binding and rapid off rate allow the Dvl proteins to interrogate various sites and search the interior cell surface (Fig. 7A). Monomeric Dvl binds to Fzd and dissociates in less than a second. Upon Wnt treatment, increased Fzd-Dvl interaction increases the binding between Fzd and Dvl and in turn increases the local concentration of Dvl (Fig 7B). The higher local concentration then drives Dvl to form oligomers on the cell membrane. The oligomerized Dvl has longer dwell time where reactions such as Dvl phosphorylation could happen together with LRP5/6 binding and phosphorylation/activation (Fig 7C). Dvl in these complexes is still dynamic and comes on and off the membrane, with smaller oligomers more dynamic than the bigger ones. Given sufficient stability of the modifications the released Dvl could also perturb downstream reactions, such as β-catenin turnover.

**Figure 7.**
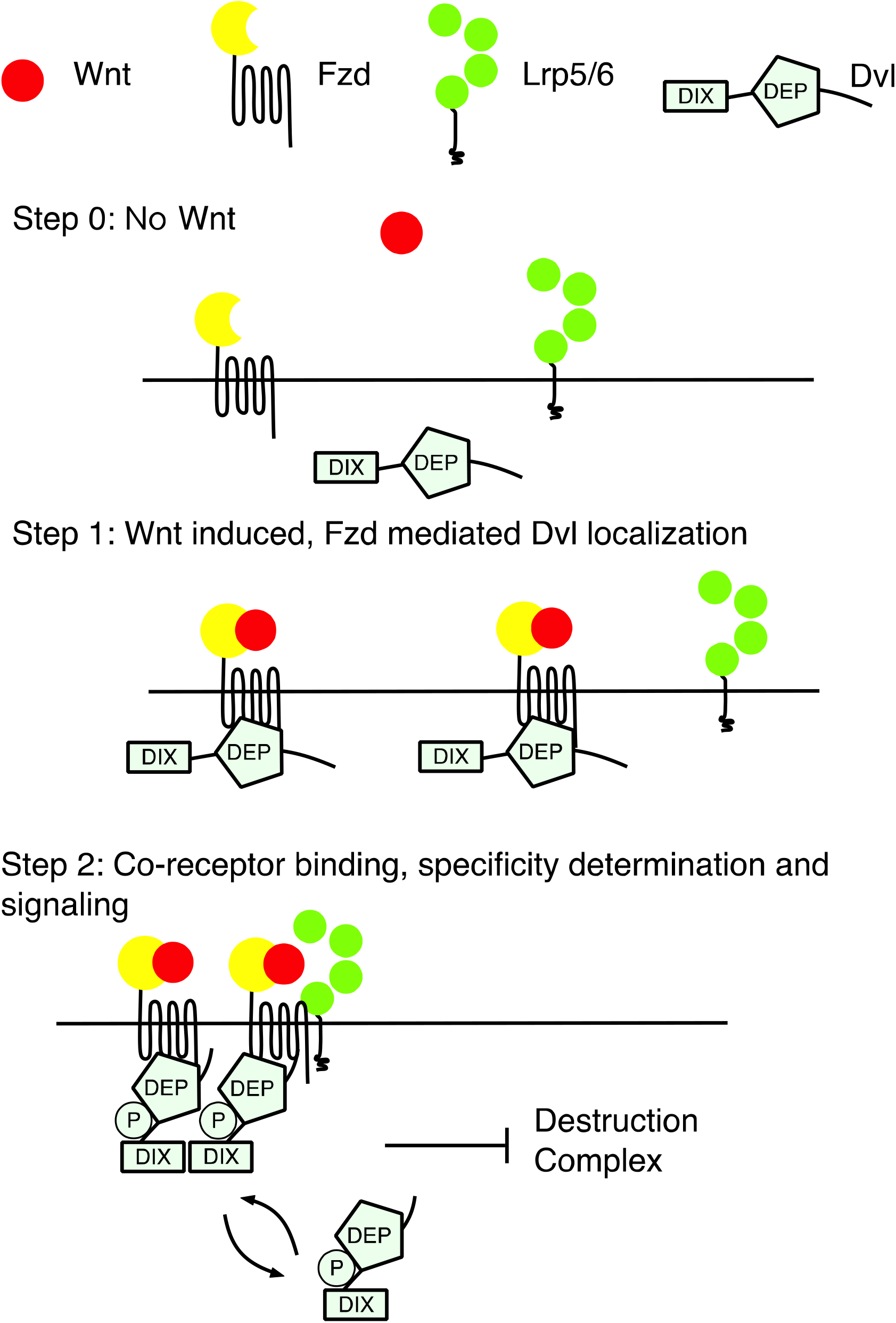
Summary of Dvl dynamics during Wnt pathway activation. (A) Majority of Dvl are monomers which binds to the membrane transiently. (B) Wnt ligands induced Dvl membrane localization which eventually leads to Dvl oligomerization. (C) Oligomerized Dvl have increased dwell time which enables potential modifications like phosphorylation to happen.

It should not be surprising that binding affinity to the membrane would in some way determine Dvl activity. It is more surprising that the effect on downstream processes reaches a maximum activity at the level of a trimer of Dvl and then declines. It is important to note a significant difference of our chimeric proteins form stable oligomers, while WT-Dvl2 forms dynamic oligomers and actively exchange the Dvl subunits in the complexes, the stable oligomer do not exchange subunits and only associate and dissociate as a whole. Furthermore, as shown in Figure 5, WT-Dvl complexes do not have a static structure. Some Dvl complexes stay longer on the membrane than expected. This must mean that the complexes can exchange subunits with the cytosolic Dvl pool. Large enough complexes can be maintained for extended period of time on the plasma membrane while still remaining inherently dynamic. Small complexes merely come on and off rapidly. We can conclude that Dvl protein complexes work best when they are both *stable* and *dynamic* on the membrane. Nature solves this seemingly incompatible task by designing Dvl protein to be a dynamic scaffold, having one membrane protein binding domain (DEP) and one oligomerization domain (DIX). The DEP domain controls the binding of Dvl to Fzd, while DIX, together with local concentration, controls the size of the Dvl oligomer. The instability of the individual Dvl subunits is related to the relative low binding affinity between DIX domains (~μM from our model) and DEP-Fzd (~μM from the experiments). When a Dvl complex gets to a certain size, it has multiple DEP domain that are incorporated into one complex, and this increases the binding to the membrane through the avidity effect. The Dvl oligomers in our study is far smaller than the “puncta” in some other studies that could reach the diameter of 1~2 um, potentially containing millions of proteins. We must point out that we didn’t rule out the possibility that other Wnt pathway proteins like Axin and APC to form large protein condensations (18) or Dvl to form super complexes in other biological systems. It’s possible that the different oligomerization forms of Dvl are used to perform different functions.

We do not yet understand why Dvl dynamicity is so carefully programmed into the system. One explanation may derive from the known and unknown phosphorylation of Dvl. Although it is possible that phosphorylation affects the dynamicity of Dvl by leading to change of binding affinities in the system. A more unusual interpretation is that the dynamicity itself may control the phosphorylation of Dvl. The rapid rate of turnover of monomeric Dvl seems to be too fast for modification by typical kinases. In this view the Wnt signal by stabilizing larger oligomers of Dvl would allow Dvl to persist long enough to be phosphorylated. The phosphorylated Dvl might then travel back into the cytosol and inhibit β-catenin degradation. Such a model would be very different from a view that Dvl, and perhaps specifically phosphorylated Dvl, only acts near the plasma membrane to inhibit the “destruction complex”. We do note however, a role of Dvl phosphorylation in this process remains enigmatic and speculative.

Another puzzling feature of the canonical Wnt signaling is the apparently loose pairings of the 19 Wnt ligands and the 10 Fzd receptors. An expected strict specificity of the ligands for one or a few receptors has not been observed. Another level of indeterminacy is how ligands and receptors activate the canonical pathway under one set of ligands leading to β-catenin stabilization and transcriptional activation and under another set of conditions activate various non-canonical pathways. Different Wnt pathways normally recruit different coreceptors but they converge at the Dvl level and diverge again thereafter. Although little is known, it is possible that differences in Dvl dynamics could play a key role in differentiating the downstream responses. Significantly we have found that the distribution of Dvl molecules are displaced to higher levels of oligomerization with the Wnt5a ligand, which activates JNK in the planar cell polarity pathway, than with the Wnt3a, a standard activator of the canonical pathway. The size distribution of membrane bound Dvl, which is nearly exponential in the canonical pathway, with Wnt5a shows a very clear peak displaced to higher molecular weight. This much less dynamic pool of Dvl oligomers on the membrane in the PCP pathway could lead to a different pattern of posttranslational modification or other interactions at the membrane. This clear difference should encourage a look at other circuits affected by the changed Dvl dynamics.

## Author contributions

W.M., M.C., X.H. and M.W.K. designed research; W.M., M.C. and H.K. performed research; Z.S. and S.A. contributed new reagents; W.M. and M.C. analyzed data; W.M. built mathematical models; W.M. and M.W.K wrote the paper.

## Acknowledgements

We thank the Nikon Imaging Center at Harvard Medical School for providing imaging environment and discuss the imaging details. We thank the Imaging and Data Analysis core facility at Harvard Medical School on providing help on analyzing the single molecule data. We specially thank Feng Cong and Xiaomo Jiang for important contributions to our work. We thank Derek Bowie and Mark Aurousseau for providing GlyRa1-GFP plasmids and discussing single molecule quantification methods. We thank Ana R. Hernández, Adrian Salic, Taran Gujral, Ren Sheng, Bryan MacDonald, Kangmin He, and Ying Lu for many helpful discussions. M.K. acknowledges support by NIGMS GM26875 and NICHD HD0731. X.H acknowledges support by R01GM057603 and R01GM126120, and the Harvard Digestive Disease Center (NIH P30DK034854) and Boston Children’s Hospital Intellectual and Developmental Disabilities Research Center (NIH P30HD18655). X.H. is an American Cancer Society Research Professor.

## Materials and methods

### Plasmids

Human Dvl2 wildtype cDNA sequence was acquired from Harvard PlasmID (HsCD00326703). The Dvl2 gene is then PCR amplified and moved onto pBABE-puro backbone (addgene #1764) using Gibson assembly. There is a 12 amino acids linker in between Dvl2 and mEGFP (GGAGGTAGTGGTGGATCTGGTGGATCAGGAGGTTCT). Dvl2 mutants was made by using Agilent QuickChange II mutagenesis kit (#200523) on the basis of Dvl2-mEGFP plasmid. For the Dvl2 oligomer mutations, all oligomer sequences are from table S1 of (36). SAM domain sequence are from *Arabidopsis thaliana* (SAM1) and *Ginkgo biloba* (SAM2) (37).

To construct donor vector for Dvl2-mEGFP-knockin, 5’ and 3’ homology arms (609bp and 840bp respectively) were PCR amplified using genomic DNA of HEK293T as the template and cloned into a vector containing mEGFP.

### Cell culture and cell lines

HEK293T and the derived cells were grown at 37 °C and 5% (vol/vol) CO_2_ in DMEM, 10% (vol/vol) FBS, and 1% antibiotics (penicillin/streptomycin).

**Dvl-TKO** cell line is the abbreviation of HEK293T cells with all the Dvl knocked-out, which is made by Dr. Stephane Angers’s lab. **Fzd-null** is the abbreviation of HEK293T cell with all Fzd knocked-out. **LRP-DKO** is the abbreviation of HEK293T cell with LRP5/6 knocked-out. HEK293T cells are from ATCC. **Dvl2-KI** cell line is the abbreviation of HEK293T cell with mEGFP knocked in on the c-terminus of Dvl2. Dvl2-1Mer are Dvl2-ΔDIX over-expressed in Dvl-TKO cells.

### Antibodies, Immunoblotting

Total cell lysates in NP40 lysis buffer (50 mM Tris, pH 7.5, 150 mM NaCl, 1mM EDTA, 1% NP40, 10% glycerol, 10mM NaF,10 mM Na_3_VO_4_, and protease inhibitor cocktail, pH=7.4) were prepared by gentle rotating at 4°C and cleared by centrifugation. Samples were run on an SDS-PAGE and transferred to Immobilon-P membrane. 0.015% digitonin in PBS supplemented with 50mM NaF and protease inhibitor cocktail were used to isolate the cytoplasmic fraction of β-catenin. Indirect immunochemistry using a secondary antibody conjugated with horseradish peroxidase was visualized using ECL reagents on LAS-3000 imager (FujiFilm). The following commercially available antibodies were used: from BD Transduction Laboratories: anti-β-catenin (610154, 1:2000); from Hybridoma Bank: anti-b-tubulin (E7, 1:5000); from Cell Signaling Technology, anti-Dvl2 (#3216, 1:1000); from Abcam, anti-GFP (ab13970, 1:1000).

### Dual luciferase assay

HEK 293T cells (ATCC# CRL-11268) were transfected using FuGENE HD (Promega, #E2312) in triplicate. Cells were plated in 24 well plates and transfected the following day with 50 ng SuperTopflash, 5 ng TK-Renilla. Dual luciferase reporter assays were performed using Dual-Luciferase® Reporter Assay System (Promega, #E1960) according to manufacturer’s instruction. Representative results are shown from one of three (or more) independent experiments.

### TIRF experiments

The TIRF experiments are done on Nikon Ti motorized inverted microscope with Perfect Focus System, 491 nm laser (50mW max power) with 50ms exposure time. Images or movies are acquired with MetaMorph software. The microscope has temperature and CO_2_ level control to minimize the environment perturbation on cells.

The cells are either grown on γ-Irradiated 35mm Glass Bottom Dishes (MatTek P35G-0.170-14-C) or on high precision microscope cover glasses (Marienfeld 0117650 lot. 33825 819). The glass was coated with 10ug/ml Fibronectin (Sigma/Aldrich F0895) in PBS for 10min before cells are plated. 100 nM IGK974 (Caymanchem 14072) was added to the cell culture medium when the cells are plated to decrease the autocrine Wnt signaling. The cells are cultured in normal DMEM and switched in to FluoroBrite (ThermoFisher A1896701) with 10% FBS and 100 nM IGK974 before imaging. The imaging experiments are normally done the next day after the cells are plated.

### Image analysis of TIRF experiments

The images from TIRF experiments were analyzed by U-Track(28) from Danuser Lab. The parameters used are 1.3 for PSF sigma, 0.05 for Alpha, and 5 for fit Window size. The single molecule intensity distributions and traces are then analyzed with homemade MATLAB codes.

To quantify single mEGFP intensity, we track the membrane protein with 20Hz imaging frequency and then extract all the traces longer than 20 steps. These traces are then analyzed with tdetector (41) for step analysis. The step sizes from all the traces are then fitted with Gaussian mixture model. The peak value of the first Gaussian function is used as the single mEGFP intensity value. The procedure can be applied to either control protein GlyRa1-mEGFP or Dvl2-mEGFP itself. Another way to estimate the single mEGFP intensity is by fitting the histograms of intensity of all the complexes in the cell. We compared the two method and could achieve similar single mEGFP intensity estimation. Step size estimation is more restricted to the quality of single molecule traces. Transient traces could greatly increase the noise of this method. The intensity histogram method is used in most of the experiments.

## References

1. Wodarz A, review of cell and developmental NR (1998) Mechanisms of Wnt signaling in development. doi:10.1146/annurev.cellbio.14.1.59.

2. Grigoryan T, Wend P, & … KA (2008) Deciphering the function of canonical Wnt signals in development and disease: conditional loss-and gain-of-function mutations of β-catenin in mice. doi:10.1101/gad.1686208.

3. Richter DJ, Fozouni P, Eisen MB, King N (2018) Gene family innovation, conservation and loss on the animal stem lineage. eLife 7. doi:10.7554/eLife.34226.

4. Foulquier S, et al. (2018) WNT Signaling in Cardiac and Vascular Disease. Pharmacological Reviews 70(1):68–141.

5. Logan C, Biol. NR (2004) The Wnt signaling pathway in development and disease. doi:10.1146/annurev.cellbio.20.010403.113126.

6. Stamos J, perspectives WW (2013) The β-catenin destruction complex.

7. Hernández AR, Klein AM, Kirschner MW (2012) Kinetic responses of β-catenin specify the sites of Wnt control. Science (New York, NY) 338(6112):1337–40.

8. MacDonald BT, Tamai K, He X (2009) Wnt/β-Catenin Signaling: Components, Mechanisms, and Diseases. Developmental Cell 17(1):9–26.

9. Kim S-E, et al. (2013) Wnt Stabilization of β-Catenin Reveals Principles for Morphogen Receptor-Scaffold Assemblies. Science 340(6134):867–870.

10. Taelman VF, et al. (2010) Wnt Signaling Requires Sequestration of Glycogen Synthase Kinase 3 inside Multivesicular Endosomes. Cell 143(7):1136–1148.

11. Bilić J, et al. (2007) Wnt Induces LRP6 Signalosomes and Promotes Dishevelled-Dependent LRP6 Phosphorylation. Science 316(5831):1619–1622.

12. Joslyn G, Richardson D, of the … WR (1993) Dimer formation by an N-terminal coiled coil in the APC protein.

13. Eubelen M, et al. (2018) A molecular mechanism for Wnt ligand-specific signaling. Science (New York, NY). doi:10.1126/science.aat1178.

14. Shibata N, Tomimoto Y, Crystallization … HT (2007) Crystallization and preliminary X-ray crystallographic studies of the axin DIX domain.

15. Schwarz-Romond T, et al. (2007) The DIX domain of Dishevelled confers Wnt signaling by dynamic polymerization. Nature Structural & Molecular Biology 14(6):484–492.

16. Sakai T, Katashima T, Matsushita T, Chung U (2016) Sol-gel transition behavior near critical concentration and connectivity. Polymer Journal 48(5):629–634.

17. Li P, et al. (2012) Phase transitions in the assembly of multi-valent signaling proteins.

18. Schaefer KN, Peifer M (2019) Wnt/Beta-catenin signaling regulation and a role for biomolecular condensates. Developmental cell 48(4):429–444.

19. Axelrod JD, Miller JR, ulman J, Moon RT, Perrimon N (1998) Differential recruitment of Dishevelled provides signaling specificity in the planar cell polarity and Wingless signaling pathways. Genes & development 12(16):2610–2622.

20. Cliffe A, Hamada F, Biology BM (2003) A role of Dishevelled in relocating Axin to the plasma membrane during wingless signaling.

21. Schwarz-Romond T, Metcalfe C, Bienz M (2007) Dynamic recruitment of axin by Dishevelled protein assemblies. Journal of Cell Science 120(14):2402–2412.

22. Bienz M (2014) Signalosome assembly by domains undergoing dynamic head-to-tail polymerization. Trends in biochemical sciences 39(10):487–495.

23. Smalley M, Signoret N, of cell … RD (2005) Dishevelled (Dvl-2) activates canonical Wnt signalling in the absence of cytoplasmic puncta. doi:10.1242/jcs.02647.

24. Lee Y-N, Gao Y, Wang H (2008) Differential mediation of the Wnt canonical pathway by mammalian Dishevelleds-1, -2, and -3. Cellular Signalling 20(2):443–452.

25. Wynshaw-Boris A (2012) Dishevelled: in vivo roles of a multifunctional gene family during development. Current topics in developmental biology 101:213–35.

26. Cervenka I, et al. (2016) Dishevelled is a NEK2 kinase substrate controlling dynamics of centrosomal linker proteins. Proceedings of the National Academy of Sciences 113(33):9304–9309.

27. Seto ES, Bellen HJ (2006) Internalization is required for proper Wingless signaling in Drosophila melanogaster. The Journal of cell biology 173(1):95–106.

28. Jaqaman K, et al. (2008) Robust single-particle tracking in live-cell time-lapse sequences. Nature Methods 5(8):nmeth.1237.

29. Fiedler M, of the … M-TC (2011) Dishevelled interacts with the DIX domain polymerization interface of Axin to interfere with its function in down-regulating β-catenin. doi:10.1073/pnas.1017063108.

30. Chiu CW, et al. (2016) SAPCD2 controls spindle orientation and asymmetric divisions by negatively regulating the G i-LGN-NuMA ternary complex. Developmental cell 36(1):50–62.

31. Gammons MV, Renko M, Johnson CM, Rutherford TJ, Bienz M (2016) Wnt Signalosome Assembly by DEP Domain Swapping of Dishevelled. Molecular cell 64(1):92–104.

32. Wong H-C, et al. (2003) Direct Binding of the PDZ Domain of Dishevelled to a Conserved Internal Sequence in the C-Terminal Region of Frizzled. Molecular Cell 12(5):1251–1260.

33. Green J, Nusse R, van Amerongen R (2014) The role of Ryk and Ror receptor tyrosine kinases in Wnt signal transduction. Cold Spring Harbor perspectives in biology 6(2). doi:10.1101/cshperspect.a009175.

34. Lu Y, Wang W, Kirschner MW (2015) Specificity of the anaphase-promoting complex: a single-molecule study. Science 348(6231):1248737.

35. Bernatík O, et al. (2014) Functional Analysis of Dishevelled-3 Phosphorylation Identifies Distinct Mechanisms Driven by Casein Kinase 1□ and Frizzled5. J Biol Chem 289(34):23520–23533.

36. Brühmann S, et al. (2017) Distinct VASP tetramers synergize in the processive elongation of individual actin filaments from clustered arrays. Proceedings of the National Academy of Sciences of the United States of America 114(29):E5815–E5824.

37. Sayou C, et al. (2016) A SAM oligomerization domain shapes the genomic binding landscape of the LEAFY transcription factor. Nature communications 7:11222.

38. Gao C, Chen Y-G (2010) Dishevelled: The hub of Wnt signaling. Cellular Signalling 22(5):717–727.

39. Wallingford JB, Habas R (2005) The developmental biology of Dishevelled: an enigmatic protein governing cell fate and cell polarity. Development 132(20):4421–4436.

40. Kirschner M, of the National GJ (1998) Evolvability. doi:10.1073/pnas.95.15.8420.

41. Chen Y, Deffenbaugh NC, Anderson CT, Hancock WO (2014) Molecular counting by photobleaching in protein complexes with many subunits: best practices and application to the cellulose synthesis complex. Molecular biology of the cell 25(22):3630–3642.

