## supplement materials for "Single molecule dynamics of Dishevelled at the plasma membrane and Wnt pathway activation"

**Methods and Material:**

***Dvl2-Crispr sequence:***

The following are the two knock-in sequences we used in the experiment. Red letters indicate the palm region of CRISPR sequence.

GAGCAAGCACATGACGGCC AGG

GTGCTTGCTCTTACAGTGCC TGG

**Text S1. Dvl membrane binding model.**

Let’s think about the simplest possible model for this system. In this model there are two populations of Dvl, one on the membrane and on in the cytosol. Dvl has basic property to form linear oligomer by the oligomerization of DIX-DIX domain interaction. It can also be recruited to the membrane primarily through Dvl-Fzd interaction. So we consider two interactions: Dvl-Dvl interaction and Dvl-Fzd interaction in the paper. The dissociation constants between monomeric Dvl-Dvl and Dvl-Fzd are defined as *K_D_* and *K_F_* respectively, by assuming ***independence between different interactions***. We use *d_n,m_* to denote ***one specific configuration*** of the Dvl complex with *n* Dvl subunits and *m* Fzd subunits. *m=0* represents the cytosolic Dvl complexes and *m>0* represents membrane bound Dvl complexes. We denote one specific configuration because Dvl complex with size n has n possible binding sites for Fzd. Fzd binding to different subunits of Dvl results in complexes with different configuration, although the affinity between Dvl and Fzd for different configurations is always *K_F_*. We have the following possible reactions:

$$d_{n-j,0}+d_{j,0}\underset{\Leftrightarrow}{K_{D}}d_{n,0}$$

$$d_{n,0}+Fzd\underset{\Leftrightarrow}{K_{F}}d_{n,1}$$

$$d_{n-j,m-k}+d_{j,k}\underset{\Leftrightarrow}{{\alpha K}_{D}}d_{n,m}$$

$$d_{n,m-1}+Fzd\underset{\Leftrightarrow}{{\alpha K}_{F}}d_{n,m} (n\geq m>1)$$

Then we use *D_n,m_* to indicate Dvl complexes with *n Dvl* and *m Fzd*, summing up all the different configurations. We then have:

$$\frac{D_{n,0}Fzd}{D_{n,1}}=\frac{K_{F}}{n}$$

$$\frac{D_{n,m}Fzd}{D_{n,m+1}}=\frac{{\left( m+1 \right)K}_{FM}}{n-m}=\frac{\alpha{\left( m+1 \right)K}_{F}}{n-m} (n-1>m>1)$$

$$\frac{D_{n-j,0}D_{j,0}}{D_{n,0}}=K_{D}$$

The equation $d_{n-j,m-k}+d_{j,k}\underset{\Leftrightarrow}{{\alpha K}_{D}}d_{n,m}$ will yield very complex relationship for *D_n-j,m-k_*, *D_j,k_* and *D_n,m_*, but we don’t need this to derive the size distribution because any complex with *n* Dvl and *m* Fzd can be imagined to first form cytosolic Dvl complex with size n, and then adding m Fzd at different site on top of the Dvl complex. We then have:

$$D_{n,m}=\left\{ \begin{aligned} {\alpha K}_{D}{\binom{n}{m}\left( \frac{Dvl}{K_{D}} \right)}^{n}\left( \frac{Fzd}{\alpha K_{F}} \right)^{m} (n\geq m>0) \\ K_{D}\left( \frac{Dvl_{N}}{K_{D}} \right)^{n} (m=0) \end{aligned} \right.$$

We can simplify the formulas as:

$$D_{n,m}=\left\{ \begin{aligned} {\alpha K}_{D}{\binom{n}{m}\delta}^{n}\left( \frac{\epsilon}{\alpha} \right)^{m} (n\geq m>0) \\ K_{D}\delta^{n} (m=0) \end{aligned} \right.$$

In turn we have the membrane Dvl molecules with n subunit as:

$$D_{n}=\sum_{m=1}^{n} D_{n,m}={\alpha K}_{D}\delta^{n}(\left( 1+\frac{\epsilon}{\alpha} \right)^{n}-1)$$

We could then write the total number of Dvl and Fzd as:

$$Dvl_{0}=\sum_{n=1}^{\infty} \sum_{m=0}^{n} nD_{n,m}=\sum_{n=1}^{\infty} nK_{D}(\alpha\delta^{n}\left( \left( 1+\frac{\epsilon}{\alpha} \right)^{n}-1 \right)+\delta^{n})=K_{D}(\frac{\alpha\delta\left( 1+\frac{\epsilon}{\alpha} \right)}{\left( 1-\delta\left( 1+\frac{\epsilon}{\alpha} \right) \right)^{2}}+\left( 1-\alpha\right)\frac{\delta}{\left( 1-\delta\right)^{2}})$$

$$Fzd_{0}=\sum_{n=1}^{\infty} \sum_{m=1}^{n} mD_{n,m}+F=\sum_{n=1}^{\infty} K_{d}\delta^{n}n\epsilon\left( 1+\frac{\epsilon}{\alpha} \right)^{n-1}+F=\frac{K_{d}\delta\epsilon}{\left( 1-\delta\left( 1+\frac{\epsilon}{\alpha} \right) \right)^{2}}+K_{F}\epsilon$$

We can solve free Dvl and free Fzd and in turn get all the following quantities.

The total Dvl complex on the membrane is:

$$D_{MC}=\sum_{n=1}^{\infty} \sum_{m=1}^{n} D_{n,m}=\sum_{n=1}^{\infty} K_{D}\alpha\delta^{n}\left( \left( 1+\frac{\epsilon}{\alpha} \right)^{n}-1 \right)=\frac{K_{D}\delta\epsilon}{(1-\delta(1+\frac{\epsilon}{\alpha}))(1-\delta)}$$

Membrane bound Dvl monomer is:

$$D_{1,1}=K_{D}\delta\epsilon$$

Total Dvl subunits on membrane is:

$$D_{M_{All}}\sum_{n=1}^{\infty} \sum_{m=1}^{n} nD_{n,m}=K_{D}\left( \frac{\alpha\delta\left( 1+\frac{\epsilon}{\alpha} \right)}{\left( 1-\delta\left( 1+\frac{\epsilon}{\alpha} \right) \right)^{2}}-\alpha\frac{\delta}{\left( 1-\delta\right)^{2}} \right)=\frac{K_{D}\delta\epsilon(1-\delta^{2}\left( 1+\frac{\epsilon}{\alpha} \right))}{\left( 1-\delta\left( 1+\frac{\epsilon}{\alpha} \right) \right)^{2}\left( 1-\delta\right)^{2}}$$

Average size of membrane bound Dvl is:

$$\bar{N}=\frac{D_{M_{All}}}{D_{MC}}=\frac{(1-\delta^{2}\left( 1+\frac{\epsilon}{\alpha} \right))}{\left( 1-\delta\left( 1+\frac{\epsilon}{\alpha} \right) \right)\left( 1-\delta\right)}=\frac{(1-\left( \frac{Dvl}{K_{D}} \right)^{2}\left( 1+\frac{Fzd}{\alpha K_{F}} \right))}{\left( 1-\frac{Dvl}{K_{D}}\left( 1+\frac{Fzd}{\alpha K_{F}} \right) \right)\left( 1-\frac{Dvl}{K_{D}} \right)}$$

**Text S2. Photobleach or simple dissociation show geometric dwell time distribution:**

Due to photobleach or complex dissociation, membrane molecule will have limited dwell time, which could be estimated using the following model.

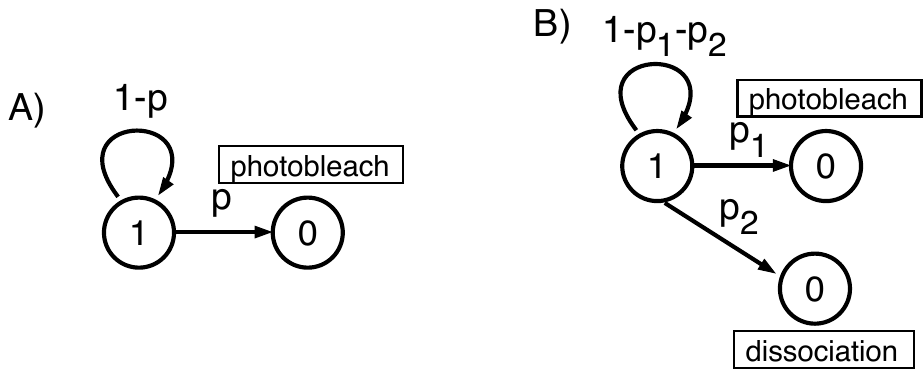

In example A), the dwell time is only affected by photobleach. The probability to find trace of length n will be $P\left( n \right)\sim p\left( 1-p \right)^{n-1}$, which is exponential distribution. Example B) doesn’t have fundamental difference. The probability to find trace of length n will be $P\left( n \right)\sim(p_{1}+p_{2})\left( 1-p_{1}-p_{2} \right)^{n-1}$. The dissociation rate can be estimated from the log probability of dwell time distribution. The slope of the log(probability) histogram is log(1-p) where $p=rate\cdot\Delta t$. Thus, we could derive the photobleach rate and dissociation rate from the dwell time distribution.

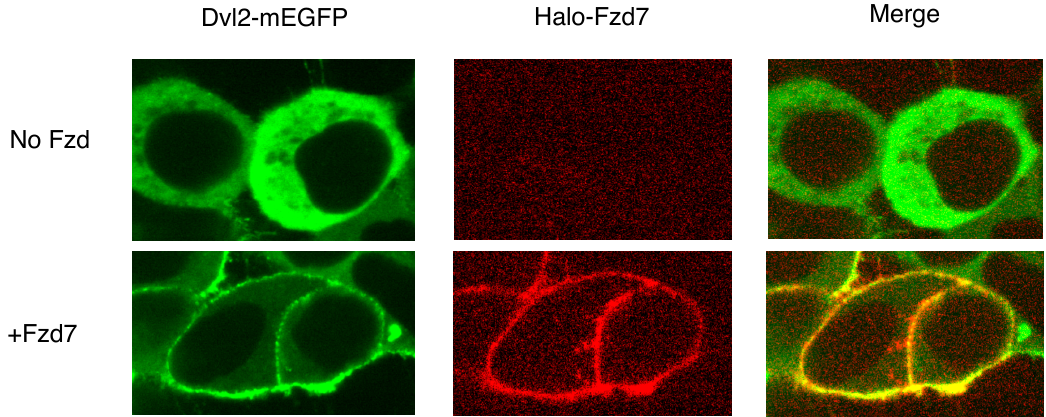

Figure S1: Dvl2 localize to the cell membrane in Fzd7 overexpressing cells.

The top panels are Dvl2-KI cells without Fzd7 over-expression. Dvl2 are in the cytosol. The bottom panels are cells with Fzd7 overexpressed. Dvl are localized to the cell membrane in these cells.

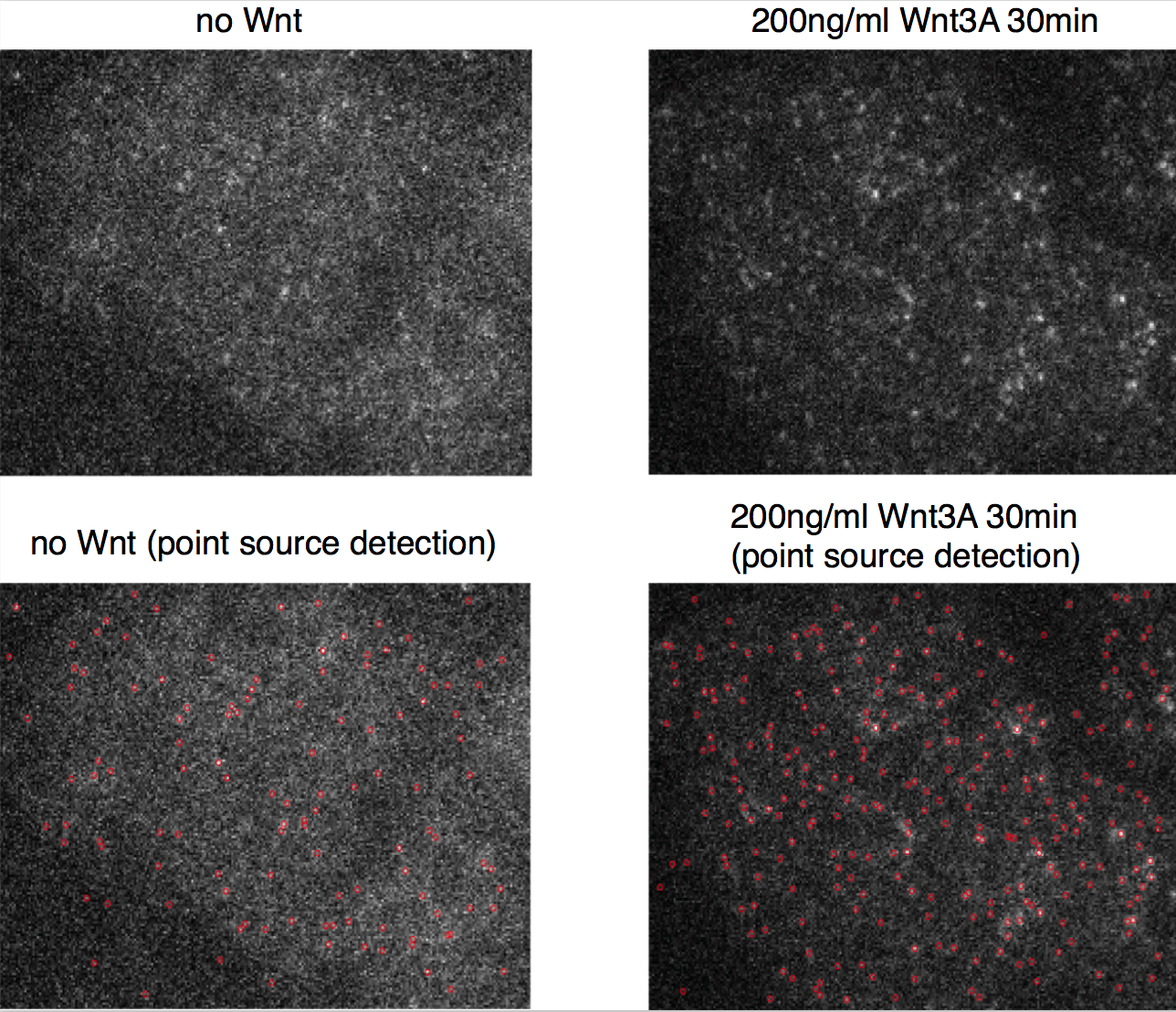

Figure S2: Examples of image analysis using uTrack.

The upper panels show the original image, the lower panels show single molecule identification results using uTrack software. Each red circle is an identified single molecule spot.

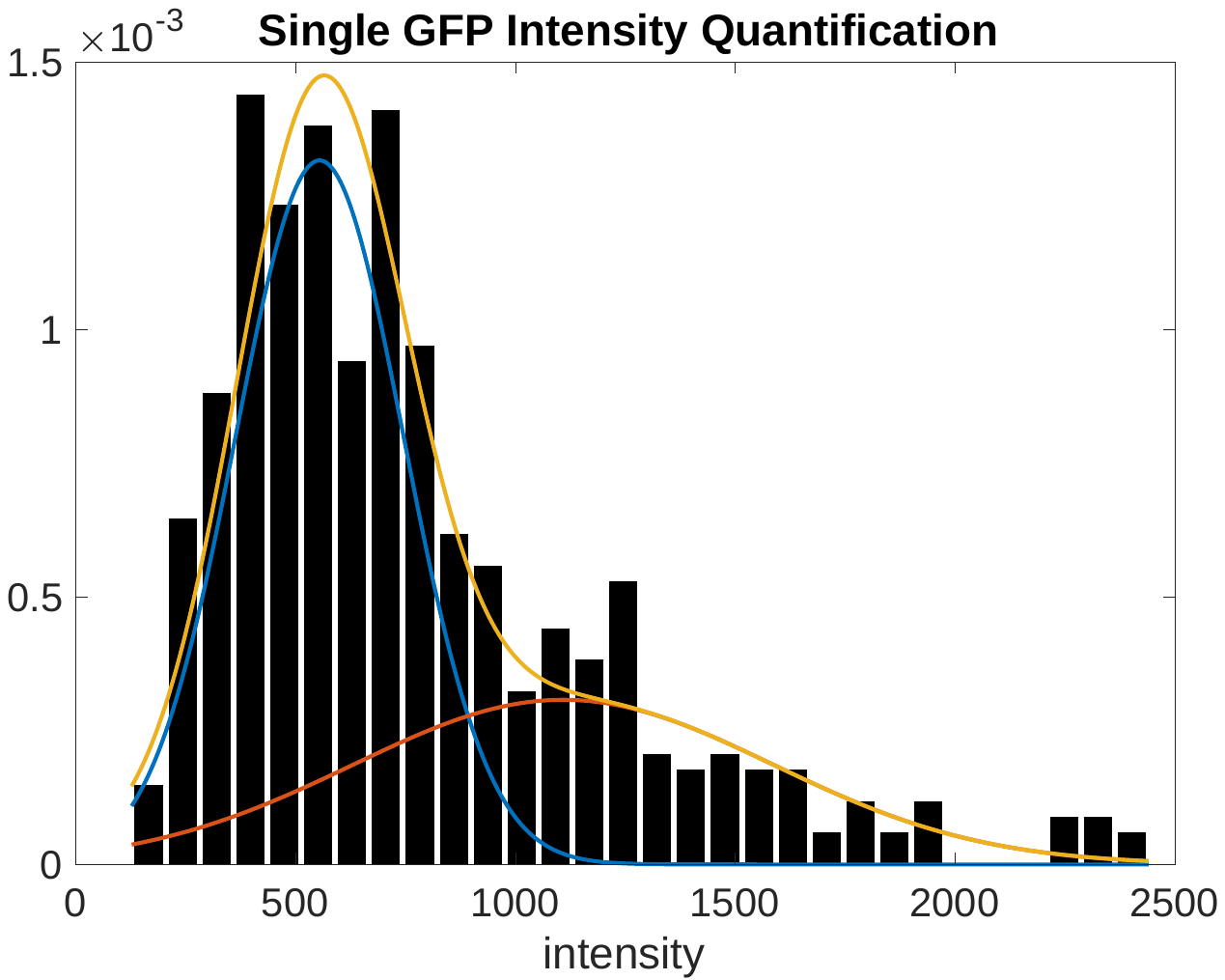

Figure S3: single GFP intensity quantification with membrane protein photobleach.

pRK5-mEGFP-GlyRa1 plasmid is used. The step jumps are fitted with two Gaussian functions and the first peak position is 554.

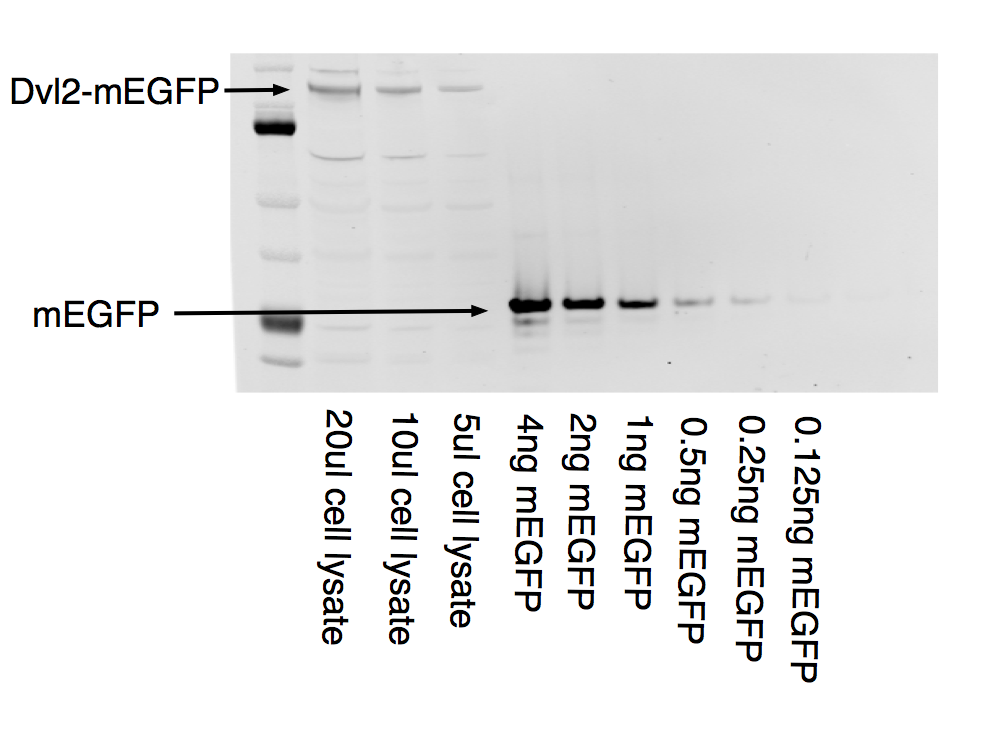

Fig S4: Quantifying Dvl2-mEGFP concentration.

1.3 million HEK293T cells with Dvl2-mEGFP was lysed to get 250ul of cell lysate. The lysate was then run together with different amount of mEGFP to estimate the concentration of mEGFP in the cell lysate. All the mEGFP lanes are loaded with 20ul sample. The concentration of the 4ng mEGFP sample is 7.07nM. The concentration of Dvl2-mEGFP in the cell lysate is then estimated to be 1.42nM. We then calculated the mean HEK293T cell volume to be 1.94pL based on the mean HEK293T cell diameter which is measured to be 15.48μm. With the cell volume and total cell number we estimated the undiluted total cell volume to be 2.54μL. Thus the actual Dvl2-mEGFP concentration is **140nM**.

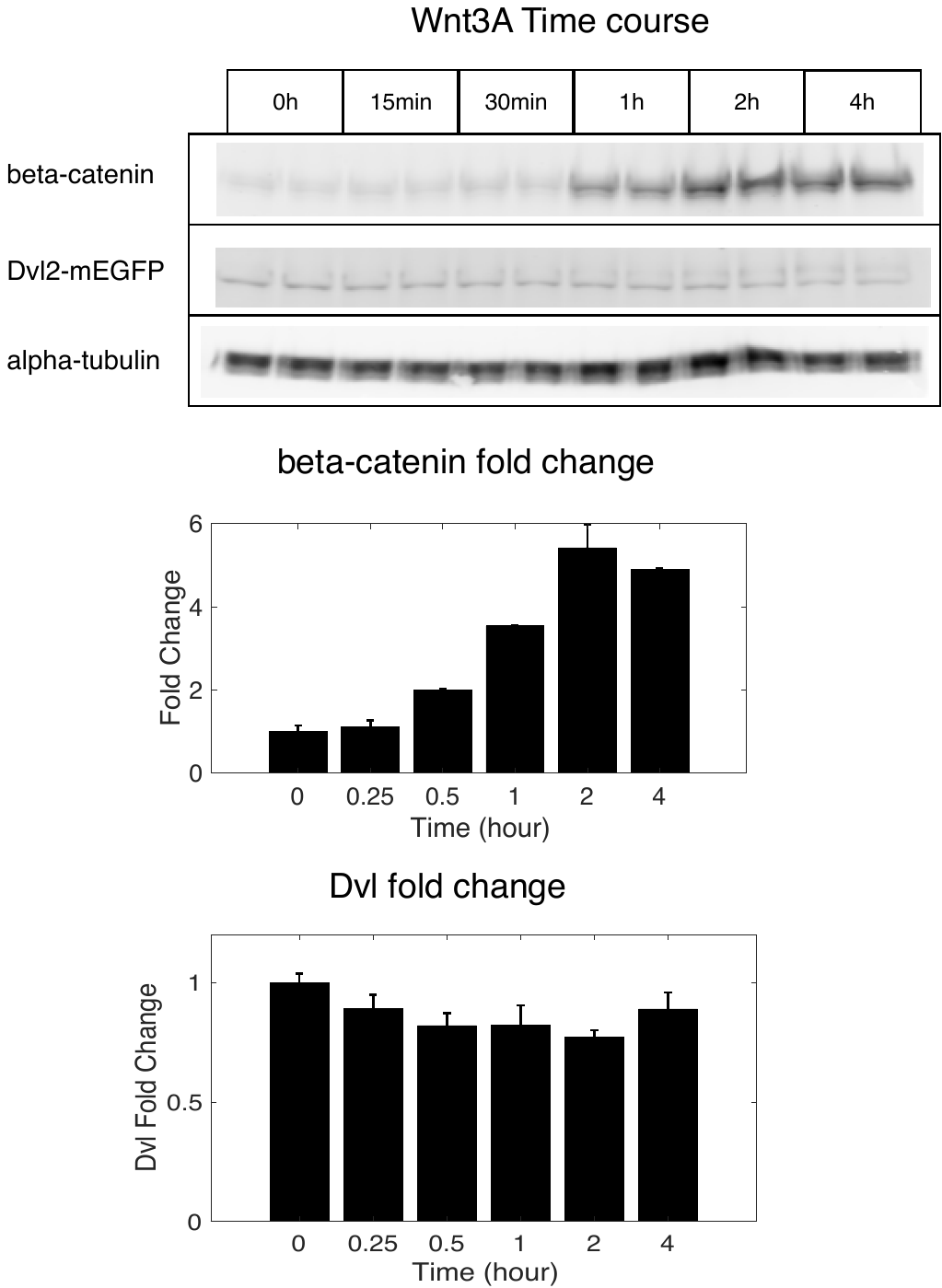

Figure S5. beta-catenin and Dvl2 level change during Wnt3A activation in Dvl2-KI cell.

Wnt3A CM is added at time 0 and the cytosolic β-catenin and Dvl2 protein levels are measured with western blot.

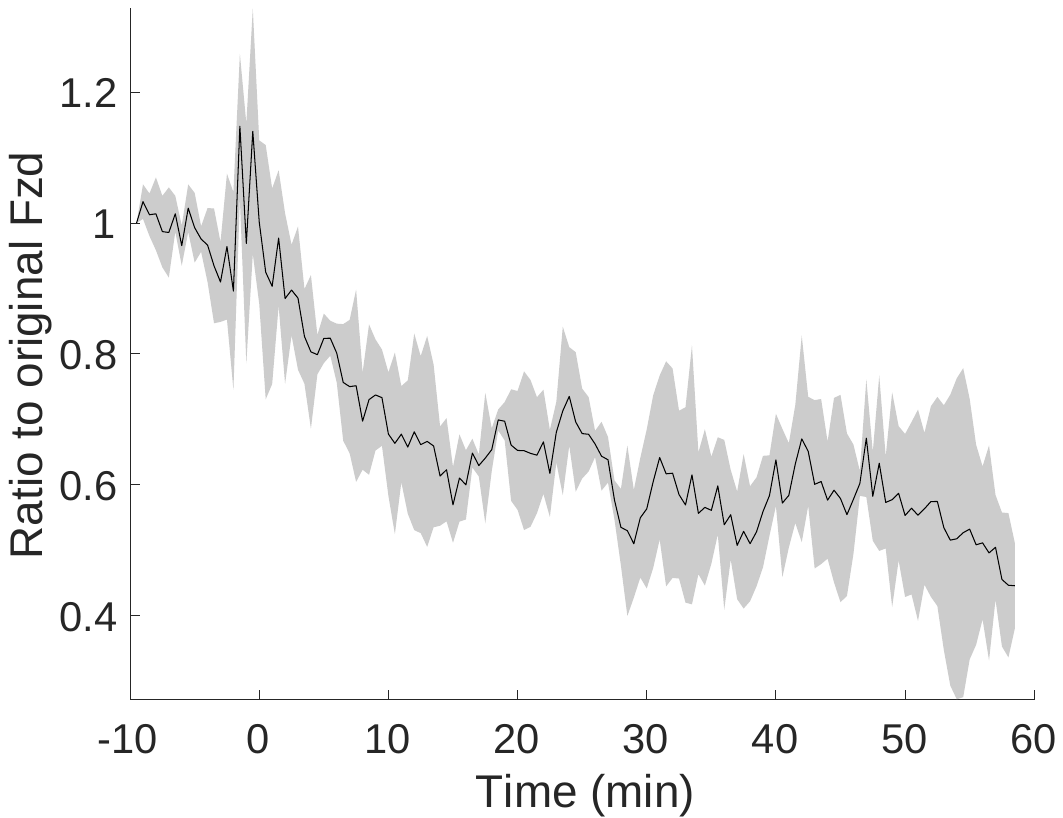

Figure S6. Ratio of membrane Fzd after Wnt3A treatment.

500ng/ml Wnt3A was added to the media at time 0. mEGFP-Fzd1 membrane total intensity is measured with TIRF microscopy and the result is the average of 3 cells.

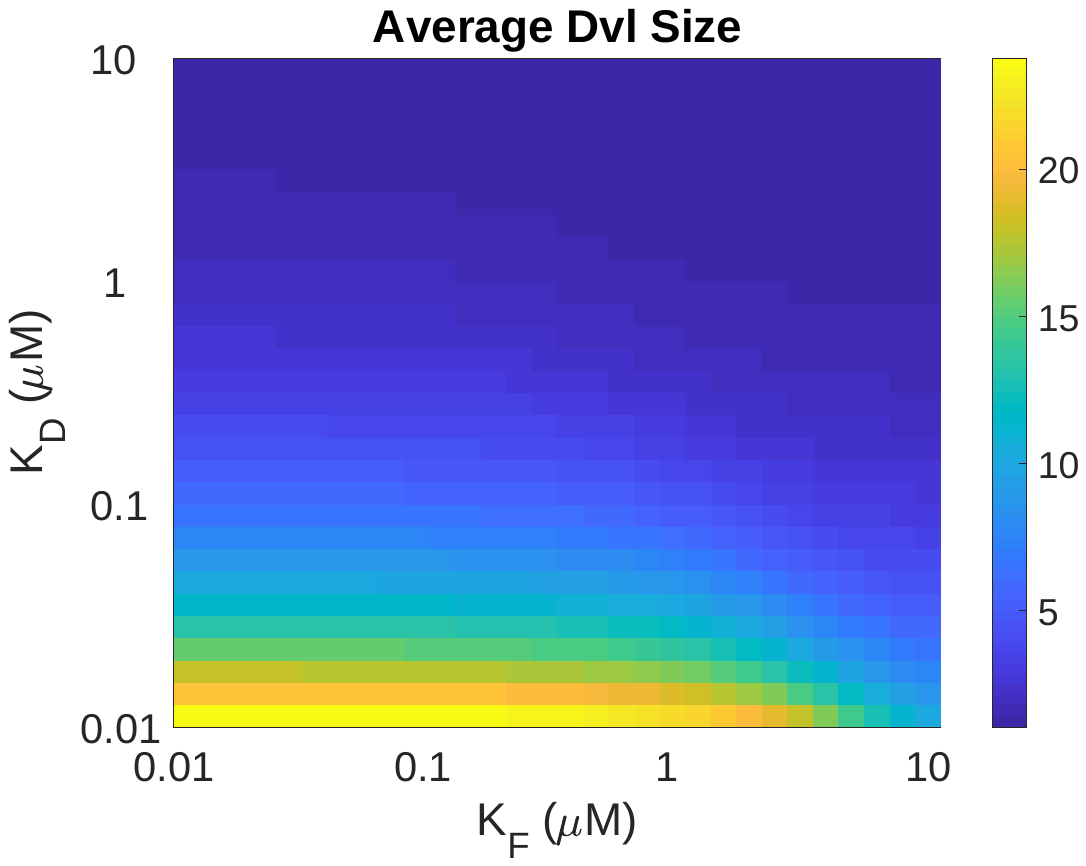

Figure S7: Model results of average membrane Dvl size varying two parameters K_D_ and K_F_. The colorbar shows the range of average size covered in the model, from 1 to 24.

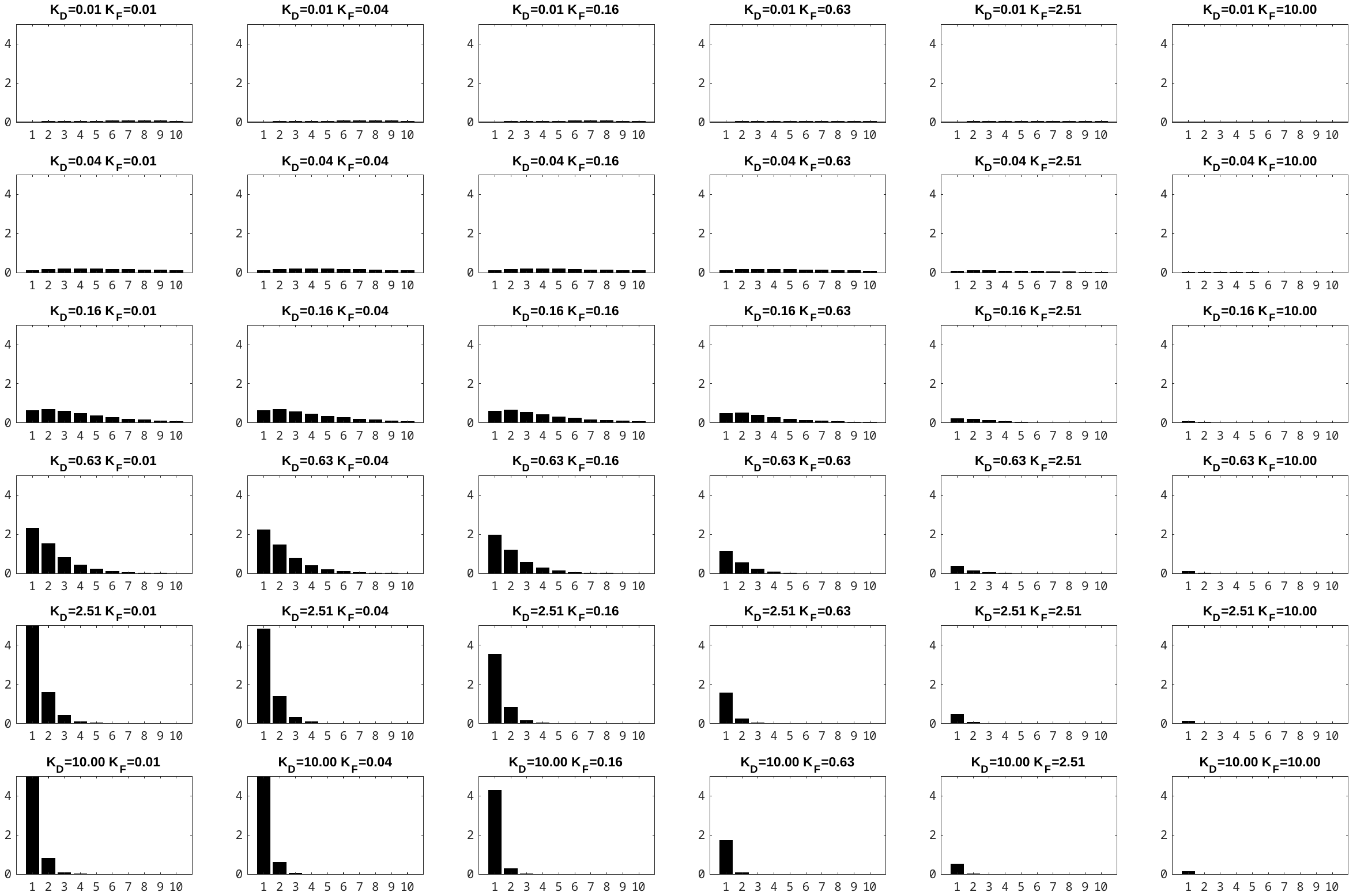

Figure S8: Model results of Dvl size distribution varying two parameters K_D_ and K_F_.

Each subfigure is a membrane Dvl size distribution. The size is on the x-axis, from 1 to 10, well the y-axis is the concentration of Dvl complexes on the cell membrane. *K_D_* is changed vertically and *K_F_* changed horizontally. The units of both *K_D_* and *K_F_* are μM. It’s easy to see that *K_D_* changes the shape of the distribution while *K­_F_* changes the absolute height of the distribution.

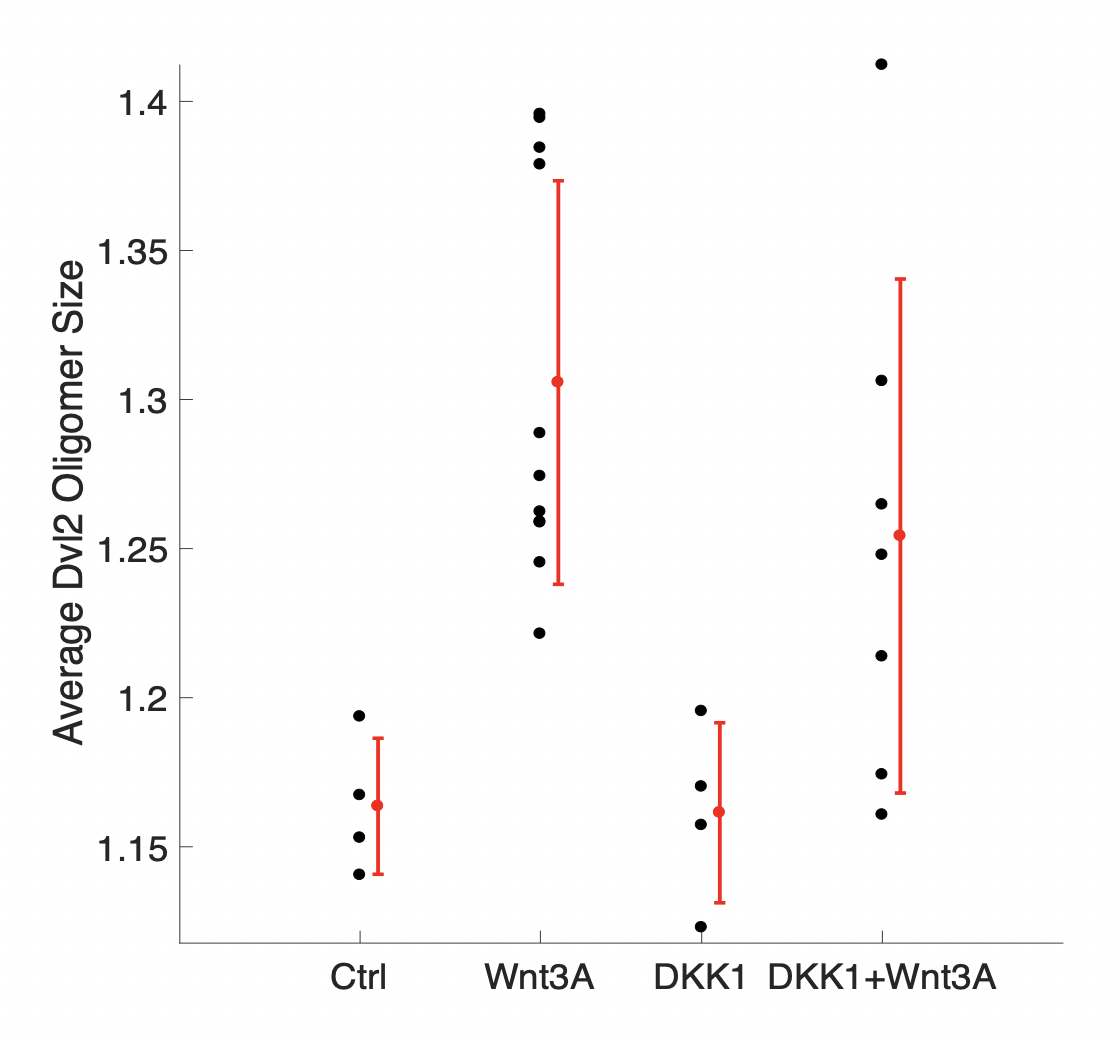

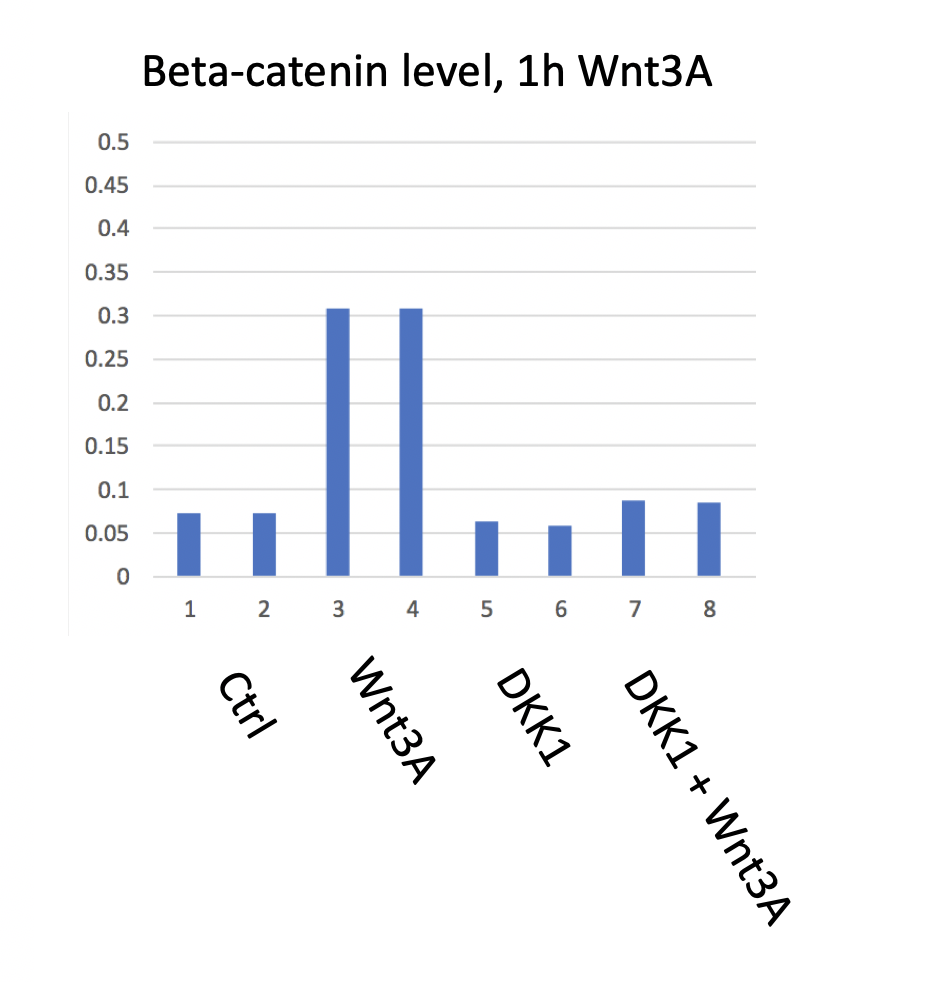

Fig S9. DKK1 blocks Wnt pathway response but not Dvl oligomerization.

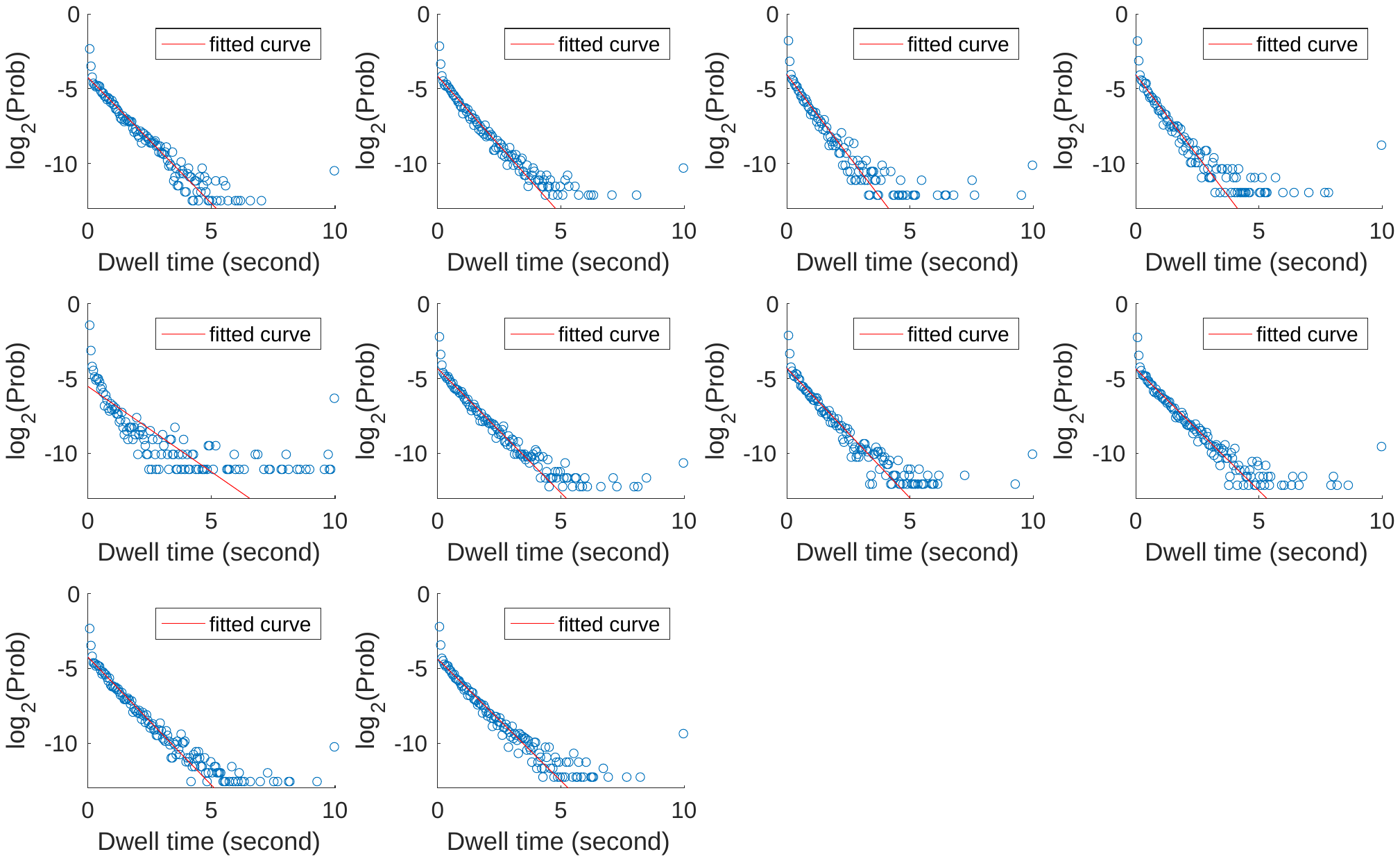

Figure S10: Dwell time distribution of membrane bound GlyRa1-mEGFP. The photobleach rate is 1.17±0.13/s, which correspond to half-life of 0.59s (11.8 frames). Each subfigure represents a different experiment, the first experiment in the second row was dropped before statistics due to poor data quality.

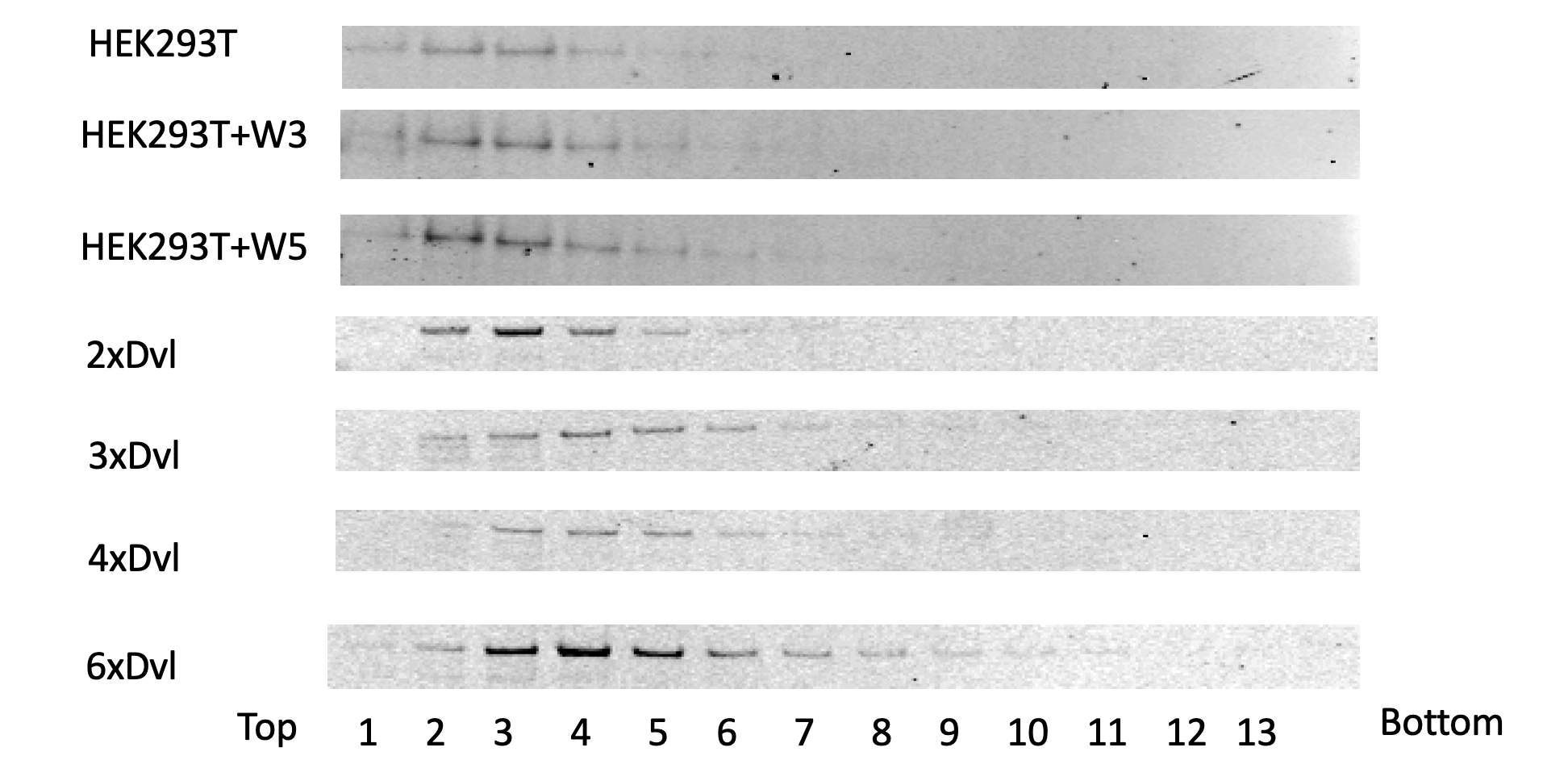

Fig S11. Sucrose gradient of cell extract. HEK293T cells are activated by Wnt3A or Wnt5A, and then broken with needle and spin down at 100,000g to remove lipids. Different Dvl mutant oligomers are overexpressed in Dvl-TKO cells and prepared the same way. The supernatants are then loaded on top of a 5~25% sucrose gradient (50mM KCl, 20mM Tris 7.5, 5mM MgCl2, 1mM EDTA, 2.5mM DTT, 0.02% sodium azide (w/v)). The gradient was span in a msl-50 rotor at 38000rpm for 8 hours. The fractions are then taken, and western blot run for Dvl2. We can see a clear shift of Dvl oligomers mutants but not from Wnt3A or Wnt5A treatment. Indicating either small population of stable Dvl complexes or weak binding between Dvl.

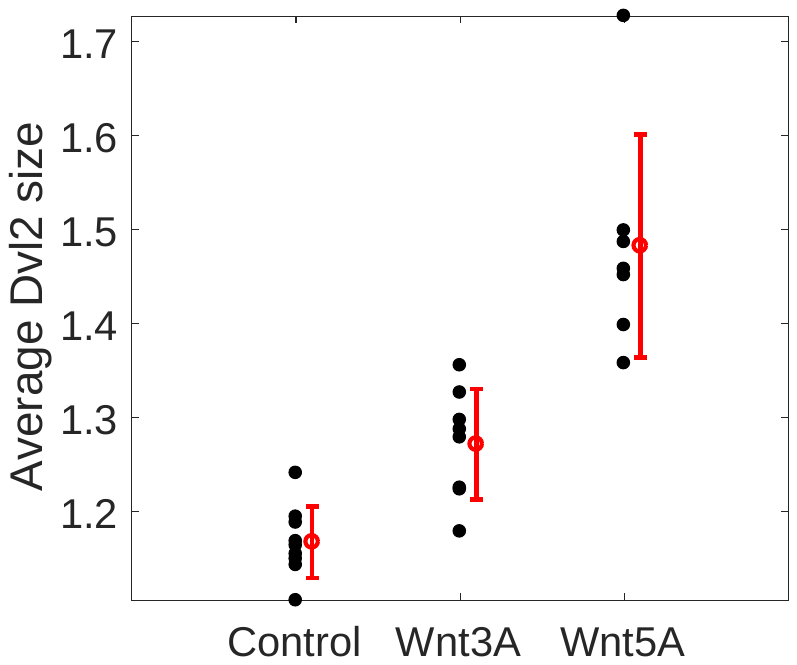

Figure S12. Average Dvl2 size under 200ng/ml Wnt3A or 200ng/ml Wnt5A treatment.

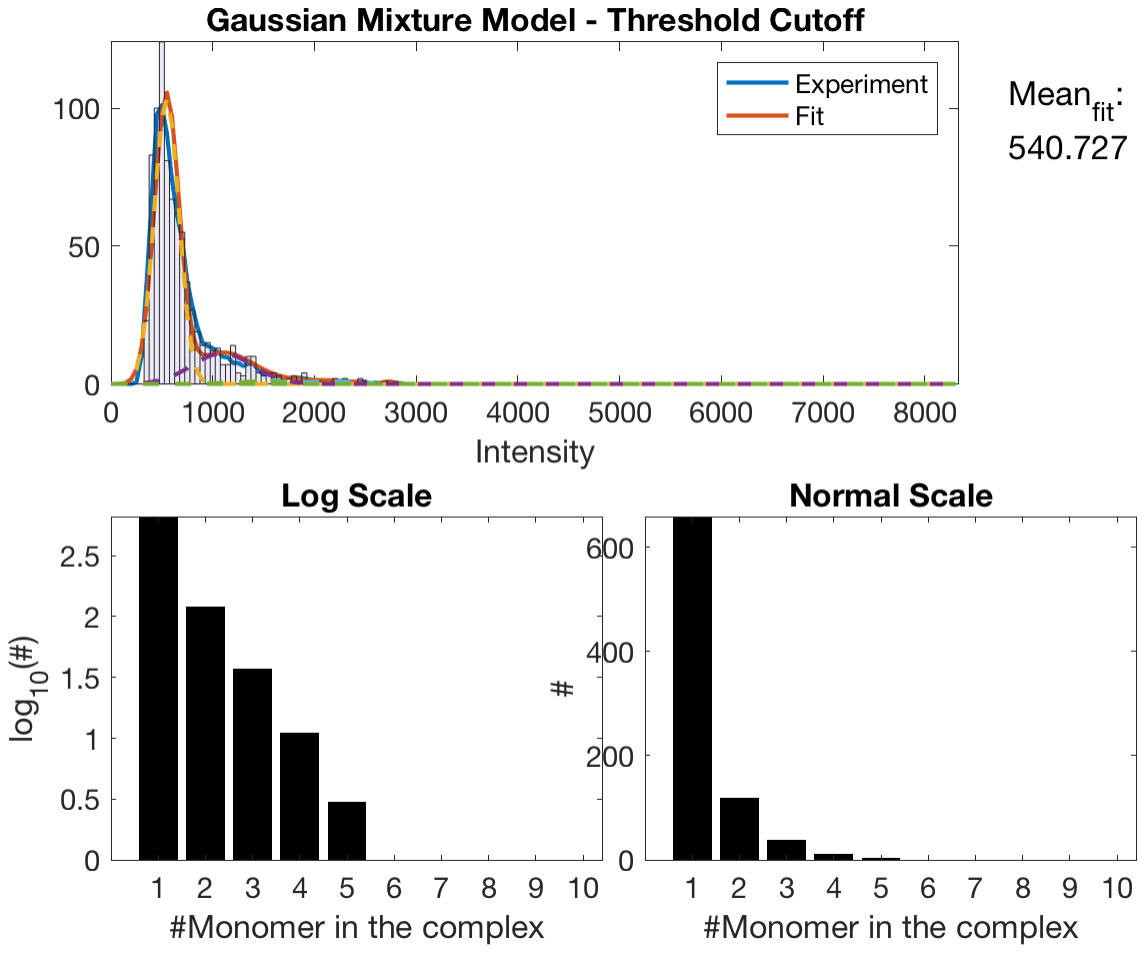

Figure S13. Typical Dvl size distribution shows exponential feature under Wnt3A treatment.

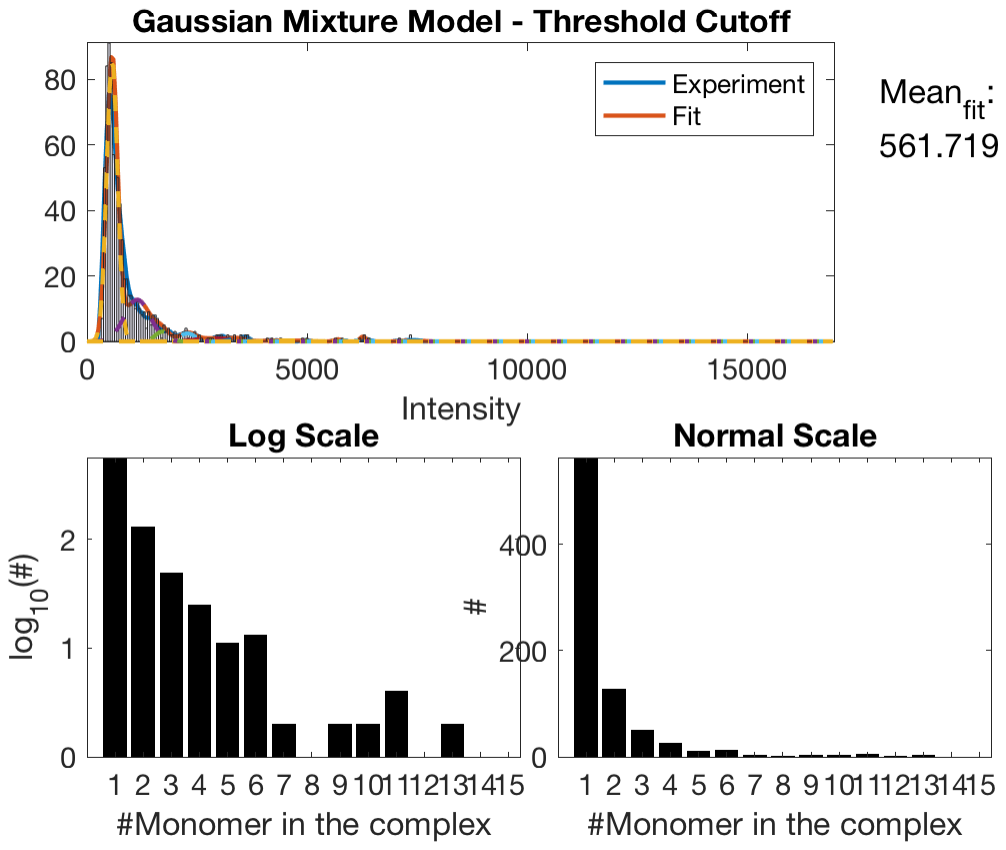

Figure S14. Typical Dvl size distribution shows long tail which can’t be explained as exponential distribution with Wnt5A treatment.
